# Deliberative Behaviors and Prefrontal-Hippocampal Coupling are Disrupted in a Rat Model of Fetal Alcohol Spectrum Disorders

**DOI:** 10.1101/2024.07.28.605480

**Authors:** Hailey L. Rosenblum, SuHyeong Kim, John J. Stout, Anna Klintsova, Amy L. Griffin

## Abstract

Fetal alcohol spectrum disorders (FASDs) are characterized by a range of physical, cognitive, and behavioral impairments. Determining how temporally specific alcohol exposure (AE) affects neural circuits is crucial to understanding the FASD phenotype. Third trimester AE can be modeled in rats by administering alcohol during the first two postnatal weeks, which damages the medial prefrontal cortex (mPFC), thalamic nucleus reuniens, and hippocampus (HPC), structures whose functional interactions are required for working memory and executive function. Therefore, we hypothesized that AE during this period would impair working memory, disrupt choice behaviors, and alter mPFC-HPC oscillatory synchrony. To test this hypothesis, we recorded local field potentials from the mPFC and dorsal HPC as AE and sham intubated (SI) rats performed a spatial working memory task in adulthood and implemented algorithms to detect vicarious trial and errors (VTEs), behaviors associated with deliberative decision-making. We found that, compared to the SI group, the AE group performed fewer VTEs and demonstrated a disturbed relationship between VTEs and choice outcomes, while spatial working memory was unimpaired. This behavioral disruption was accompanied by alterations to mPFC and HPC oscillatory activity in the theta and beta bands, respectively, and a reduced prevalence of mPFC-HPC synchronous events. When trained on multiple behavioral variables, a machine learning algorithm could accurately predict whether rats were in the AE or SI group, thus characterizing a potential phenotype following third trimester AE. Together, these findings indicate that third trimester AE disrupts mPFC-HPC oscillatory interactions and choice behaviors.

**Significance statement:** Fetal alcohol spectrum disorders (FASDs) occur at an alarmingly high rate worldwide. Prenatal alcohol exposure leads to significant perturbations in brain circuitry that are accompanied by cognitive deficits, including disrupted executive functioning and working memory. These deficits stem from structural changes within several key brain regions including the prefrontal cortex, thalamic nucleus reuniens, and hippocampus. To better understand the cognitive deficits observed in FASD patients, we employed a rodent model of alcohol exposure during the third trimester, a period when these regions are especially vulnerable to alcohol-induced damage. We show that alcohol exposure disrupts choice behaviors and prefrontal-hippocampal functional connectivity during a working memory task, identifying the prefrontal-hippocampal network as a potential therapeutic target in FASD treatment.

## Introduction

Fetal alcohol spectrum disorders (FASDs) are the most common preventable cause of developmental disability globally and are characterized by a range of physical defects and cognitive and behavioral impairments, the extent of which are dependent on the timing of exposure to alcohol (AE) (Coles, 1994; Hoyme et al., 2016; Mattson et al., 2019; Popova et al., 2023; Rasmussen, 2006). AE during the brain growth spurt, which occurs during the third trimester in humans and the first two postnatal weeks in rats (Dobbing & Sands, 1979), results in executive functioning deficits (Gursky et al., 2021; Thomas et al., 1996), which are a hallmark of FASD (Mattson et al., 2019; Rasmussen, 2006).

The medial prefrontal cortex (mPFC), hippocampus (HPC), and their interaction are important for memory-guided decision-making and are damaged after AE during the brain growth spurt (Bonthius & West, 1991; Churchwell & Kesner, 2011; Floresco et al., 1997; Ikonomidou et al., 2000; Hamilton et al., 2010, 2017; Livy et al., 2003; Lawrence et al., 2012; Maharjan et al., 2018; Murawski et al., 2012; Otero et al., 2012; Tran & Kelly, 2003; G.-W. Wang & Cai, 2006; Whitcher & Klintsova, 2008). The thalamic nucleus reuniens mediates mPFC-HPC interactions during spatial working memory (Hallock et al., 2016) and is also damaged after AE (Gursky et al., 2019, 2020), leading us to predict AE during this period would impair spatial working memory.

The HPC, mPFC, and nucleus reuniens are implicated in choice behaviors known as vicarious trial and errors (VTEs), which are thought to reflect deliberation and occur when rats pause and alternate head movements towards choice options during decision-making (Bett et al., 2012; Blumenthal et al., 2011; Griesbach et al., 1998; Hu & Amsel, 1995; Papale et al., 2012; Kidder et al., 2021; Redish, 2016; Schmidt et al., 2019; Stout et al., 2022; Tolman, 1939). VTEs emerge when flexible decision-making strategies are favored, such as when task rules are switched, and diminish with increasing task proficiency (Amemiya & Redish, 2016; Blumenthal et al., 2011; Griesbach et al., 1998; Hu & Amsel, 1995; Papale et al., 2012; Redish, 2016; Steiner & Redish, 2012). HPC lesions or disruption (Bett et al., 2012; Blumenthal et al., 2011; Griesbach et al., 1998; Hu & Amsel, 1995) and mPFC disruption (Kidder et al., 2021; Schmidt et al., 2019) result in VTE reductions. Furthermore, nucleus reuniens inactivation increases VTEs during consecutive choice error sequences, suggesting its importance for successful deliberation (Stout et al., 2022). Consequently, we predicted VTE behaviors would be disrupted after AE.

As rats approach choice points, HPC ensembles alternate between representations of potential choice trajectories ahead of the rat (Johnson & Redish, 2007, Kay et al., 2020; Tang et al., 2021). The mPFC is hypothesized to evaluate these trajectories (Redish, 2016; J. X. Wang et al., 2015), which aligns with PFC involvement in goal-directed and flexible behaviors (Miller & Cohen, 2001) and the increase in mPFC-HPC oscillatory synchrony via theta rhythms (6-10 Hz oscillations in the local field potential; LFP) during decision-making (Benchenane et al., 2010; Hallock et al., 2016; Jones & Wilson, 2005; O’Neill et al., 2013). The nucleus reuniens has been shown to transfer trajectory-relevant information from mPFC to HPC (Ito et al., 2015) and its inactivation reduces mPFC-HPC theta coherence (Hallock et al., 2016; Stout et al., 2022), suggesting a critical role in mPFC-HPC interactions. Therefore, we predicted that AE would lead to altered mPFC-HPC oscillatory activity during deliberation.

Our results show that AE during the brain growth spurt led to fewer VTEs in adulthood and resulted in a dissociation between VTEs and subsequent task performance. Despite these disruptions, task choice accuracy was unimpaired. We also demonstrate that mPFC-HPC physiology and functional connectivity were disrupted in the AE group. Lastly, we show that a machine learning algorithm could predict whether rats belonged to the AE or sham intubated (SI) group based on select behavioral measures, therefore modeling a phenotype for third trimester AE.

## Methods

### Animal subjects

Subjects were Long Evans hooded rats (5 AE female, 6 AE male; 2 SI female, 5 SI male). Choice accuracy over sessions analysis included an additional cohort of rats (9 AE female, 8 AE male; 9 SI female, 13 SI male). Pregnant dams were obtained from Charles River (Wilmington, MA). Subjects were generated from 10 litters and were born at the University of Delaware. The animal colony room was temperature and humidity controlled and followed a light/dark cycle from 7 a.m.- 7 p.m. Rats had *ad libitum* access to food and water until pretraining, when they were placed on mild food restriction to maintain 90% of their original body weight. All animal procedures followed the University of Delaware Institutional Animal Care and Use Committee (Animal Use Protocol #1177) and the NIH Guide for the Care and Use of Laboratory Animals. See Figure 1A for the experimental timeline.

**Figure 1.**
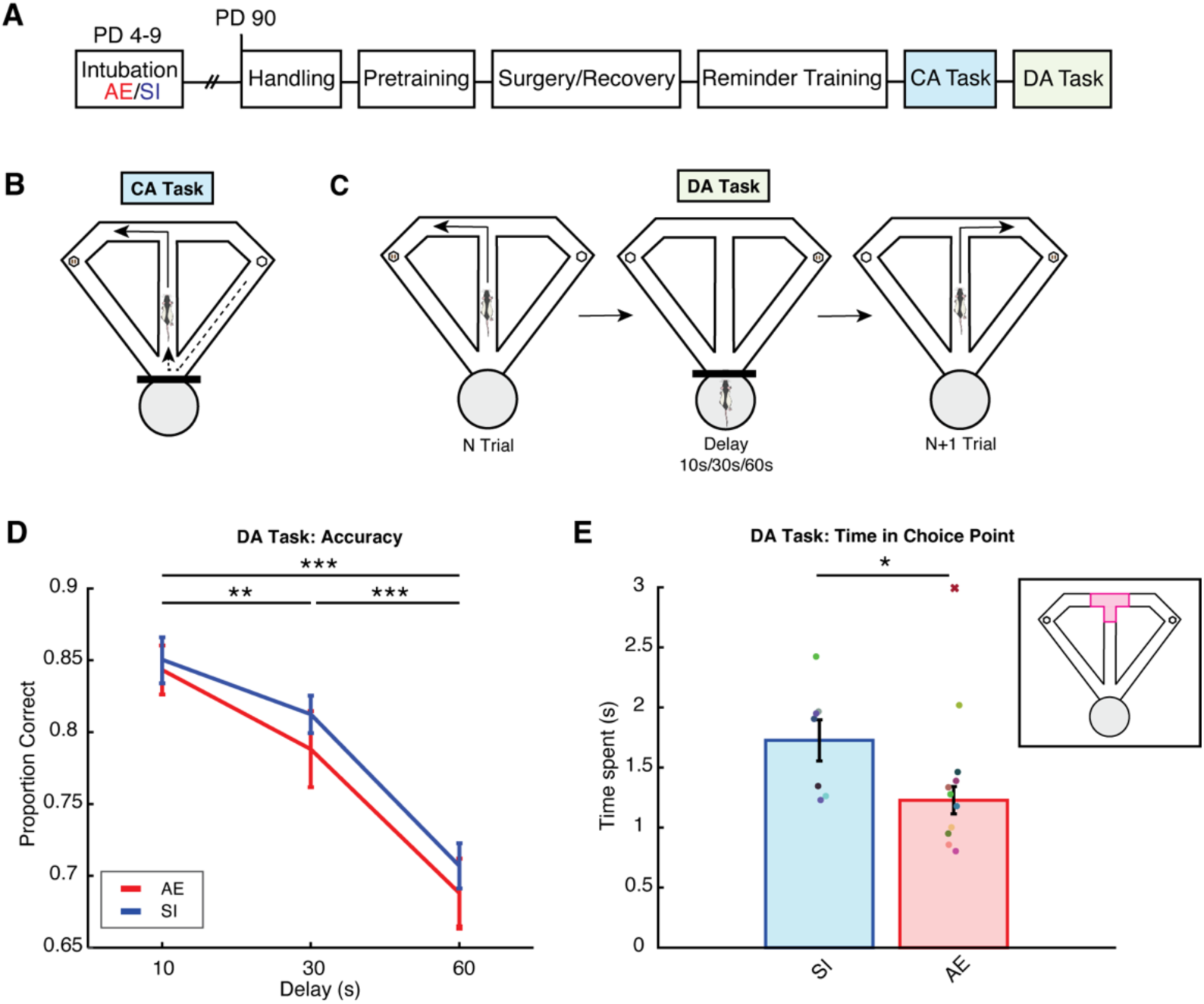
The alcohol exposed group spends less time in the choice point than the sham intubated group. **A)** Experimental timeline. PD=postnatal day, AE=alcohol exposed, SI=sham intubated, CA=continuous alternation, DA=delayed alternation. **B)** CA task schematic. Rats alternated between left and right choices over trials to receive a reward. **C)** DA task schematic. After each trial, rats returned to the start box (gray circle) to complete a delay of either 10, 30, or 60 seconds (s). **D)** DA task choice accuracy for all 10-, 30-, and 60-second delay trials in the AE (red) and SI (blue) groups. The proportion of correct trials decreases with delay length in the SI and AE groups and is not different between groups. **E)** Rats in the AE group (red) spend significantly less time in the choice point compared to rats in the SI group (blue) during the DA task. Colored dots indicate individual rats. An outlier rat in the AE group is indicated with a red “X”. Inset: T-maze with the choice point highlighted in pink. *p<0.05, **p<0.01, ***p<0.001. Error bars represent mean +/- standard error of the mean.

### Animal generation and postnatal treatment

Pups were paw marked on postnatal day 3 with an injection of India black ink and were randomly assigned to the AE or SI group. On postnatal days 4-9, pups in the AE group were administered 5.25 g/kg/day ethanol in a milk formula via intragastric intubation (divided between 2 doses at 9 a.m. and 11 a.m.). This procedure has been shown to result in a peak Blood Alcohol Concentration (BAC) of about 350 mg/dL (high dose) (Gursky et al., 2019, 2020, 2021) when measured 2 hours after the second alcohol intubation. SI pups were intubated without any liquid to control for the stress effects of intubation. To prevent weight loss, AE pups received a supplemental dose of milk formula 2 hours after the second intubation on postnatal days 4-9 and an additional dose 4 hours after the second intubation on postnatal day 4. Rats were ear punched for identification on postnatal day 9. All rats were housed with their dams until postnatal day 23, when they were weaned and pair housed until surgery.

### Behavior apparatus and testing room

Tasks were performed in a wooden T maze, which consisted of a central arm (116 cm × 10 cm), two goal arms (56.5 cm × 10 cm), and two return arms (112 cm × 10 cm) with 6 cm high wooden walls. Small weighing boats were attached at the end of each goal arm for food reward delivery. The start box at the base of the maze consisted of a barstool with a dish attached on top. Visual cues were attached to a black curtain that surrounded the room, which was dimly lit by 2 compact fluorescent bulbs.

### Handling

After postnatal day 90, experimenters handled rats for 10 minutes/day for 5 days. After each session, chocolate sprinkles were placed in the home cage to familiarize rats with the food reward of the behavioral tasks.

### Surgical procedures

Rats were anesthetized with isoflurane (1-3.5% in oxygen) and injected with atropine (0.06 mg/mL). Eye ointment was applied to the eyes and was reapplied periodically throughout the surgery. Once the pedal reflex was not displayed, their head was shaved and they were placed into a stereotaxic instrument (Kopf). The incision site was sterilized with chlorhexidine solution and injected with lidocaine. Hydrogen peroxide was used to control bleeding after the incision. After the skull was leveled and bregma was identified, a stereotaxically mounted drill was used to mark craniotomy coordinates for dorsal HPC and mPFC. Craniotomies were +3.1 mm anterior and +1.0 mm lateral to bregma (targeting prelimbic cortex) and −3.7 mm posterior and +2.2 mm lateral to bregma (targeting dorsal CA1). A cerebellum reference drill hole was made 12 mm posterior and −2.2 mm lateral to bregma. 4 bone screws (Fine Science Tools) were inserted for stability and an additional bone screw was inserted above the cerebellum for grounding. The mPFC wire bundle (2 stainless steel wires; wire diameter: 0.2 mm) was implanted 2.6 mm ventrally at an 8-degree angle. A bundle of 4 wires (each wire staggered by 0.25 mm) was implanted 2.5 mm ventrally at the HPC coordinates. The cerebellum reference wires (2 wires twisted together) were implanted 1 mm ventrally. Wires were stabilized to the skull with Metabond. Dental acrylic (Lang Dental) was used to secure a rod attached to an electrode interface board to the skull and to stabilize the wire bundles. A copper mesh cage was placed around the drive components, and a wire attached to the grounding screw was soldered to the cage and linked to the electrode interface board with a gold pin. All other wires were also linked to the electrode interface board and liquid electrical tape was applied over exposed wire. To protect drive components, a small weighing boat was velcroed on top of the copper mesh cage and the implant was wrapped in a self-adhesive bandage. Neosporin and lidocaine were applied to the skin surrounding the copper mesh. At the end of surgery, rats were injected with flunixin (Banamine; 50 mg/mL) for post-surgery analgesia. In addition, 25 mL child’s ibuprofen (100 mg/5 dL) was added to the drinking water in the home cage. Rats completed a minimum of 1 week of recovery before starting pre-training.

### Pre-training

During goal box training, rats were trained to eat chocolate sprinkles from the weighing boats in the goal zones of the maze. Wooden barriers were placed on both sides of the goal zone. Over 6 alternating trials, rats were placed in the left or right zone until they ate all the sprinkles or 3 minutes had passed. Rats were required to eat all the sprinkles in under 90 seconds during each trial over two consecutive days.

Forced run training familiarized rats with the T-maze route. Wooden barriers blocked the entry to the stem of the maze and either the left or right goal arm at the start of each trial. Once the barrier at the start box was lifted, rats traveled down the stem of the maze to the T-intersection and then proceeded down the unblocked goal arm. Rats ate the reward in the goal zone and returned to the start box via the return arm. A wooden barrier was then placed at the entry to the maze. Each session consisted of 12 trials (6 left and right in a random order). Rats spent 3-5 sessions completing the task until they performed trials without guidance from the experimenter. Before continuing training, rats were acclimated to performing the task while plugged in to the recording headstage.

### Experimental design for behavioral tasks

The continuous alternation (CA) task is an HPC-independent task (Ainge et al., 2007) that follows a spatial alternation rule (Figure 1B). To receive a reward, rats alternated between the left and right goal arms over trials without returning to the start box. Rats were required to reach a criterion of 80% choice accuracy (at least 32/40 trials correct) for two consecutive sessions.

Rats then began testing on the HPC-dependent delayed alternation (DA) task (Ainge et al., 2007; Figure 1C). Rats were rewarded for alternating left and right goal arms over trials and returned to the start box between trials to complete a delay. We systematically altered working memory load by changing the delay duration between trials (10, 30, or 60 seconds). Each DA task session consisted of 36 delay trials (plus an initial trial that rewarded rats for choosing either arm), with 12 trials of each delay length pseudorandomly interleaved within the session. LFPs were recorded from the mPFC and HPC during the task. Rats completed between 9-23 recording sessions.

### Perfusion and histology

Rats were anesthetized with isoflurane and were intraperitoneally injected with a veterinarian- approved mixture of xylazine and ketamine. Once rats no longer displayed the pedal and blink reflexes, they were transcardially perfused with 100 mL of heparinized 0.1 M phosphate buffered saline (PBS) followed by 100 mL of 4% paraformaldehyde in 0.1 M PBS (pH= 7.20). After the head was postfixed in 4% paraformaldehyde solution for 48 hours, the brain was extracted and transferred through 3 solutions of 30% sucrose in 4% formaldehyde (24-72 hours in each solution until the brain sank) and stored at 4°C until cryosectioning. A Leica cryostat (−20°C) was used to section brains in the coronal plane at 40 µm and sections were stored in rostro-caudal order in a sucrose/ethylene glycol cryoprotectant solution at - 20°C to verify electrode position. Electrode placement was verified by superimposing coronal section images on a plate from the Paxinos and Watson (2006) stereotaxic atlas.

### Video tracking and electrophysiology recordings

Video tracking data were obtained with a camera mounted to the ceiling that recorded LED lights attached to the rat’s headstage at 30 Hz (Cheetah). Video tracking data from the DA task were visually examined. Trials were excluded from analysis if they contained >10% tracking error in the stem entry to choice point exit portion of the maze or had a failed stem entry/choice point exit (i.e. video tracking lost the rat at these locations). If a trial contained a failed start box entry (when the rat returned for a delay at the end of a trial), the following trial was removed.

A 64-channel digital recording system (Digital Lynx; Neuralynx) was used to record mPFC and HPC LFPs, which were sampled at 2 kHz and filtered between 1-600 Hz using Cheetah software (Neuralynx). LFPs were examined for artifacts and corresponding trials were excluded from analysis.

### Behavioral analysis

#### Separating trials by delay length

11 AE and 7 SI rats were implanted with recording drives with LEDs on the headstage for video tracking. To examine the effect of delay length on choice accuracy and VTEs, video tracking data were used to calculate the time spent in the start box between trials. Trials were excluded from analysis if rats did not leave the start box before the start of the following delay interval (e.g., a 10-second delay trial where a rat did not exit the start box until after an actual delay of 30 seconds or greater had passed). Any 60-second delay trial with an actual delay above 100 seconds was excluded. Trials initiated more than 5 seconds before the intended delay time were also excluded (e.g., a 60-second delay trial where the trial was initiated early, and the actual delay was less than 55 seconds). This step accounted for potential disturbances in the testing room, such as the drive unplugging.

#### VTE trial identification

VTEs were identified using the integrated absolute change in angular velocity (IdPhi), a metric that captures head movement complexity (Papale et al., 2012). Low IdPhi scores reflect direct paths through the maze, whereas high IdPhi scores reflect pausing, reorienting, and head-sweeping behaviors characteristic of VTEs. First, x and y position data from the stem to the choice point exit of the maze were smoothed (*smoothdata.m*) using a moving average with a gaussian window (window size= 30; 1 second of data). A discrete-time adaptive windowing method was used to calculate velocity in the x and y dimensions (Janabi-Sharifi et al., 2000). The arctangent of the dX and dY components was taken and unwrapped to determine the orientation of motion, Phi. The change in orientation, dPhi, was calculated by applying the discrete-time adaptive windowing method to Phi. The integral of the absolute change in orientation (|dPhi|) was calculated to obtain an IdPhi score for each trial. The natural log of IdPhi was taken and lnIdPhi scores were z scored by rat. zlnIdPhi scores from AE and SI rats’ trials were shuffled before examination of the data to blind the experimenter to group.

The VTE threshold is the value where the distribution of IdPhi scores deviates from a normal distribution; this can be visualized as a “tail” off the right side of the distribution (Redish 2016, Figure 2A). Trials with scores above this threshold typically represent VTE trials, whereas scores below this threshold typically represent non-VTE trials (an example non-VTE trial is shown in the inset of Figure 2A). As the deflection point occurred at a zlnIdPhi of 0.3, this value was selected as the VTE threshold, which is similar to previously reported thresholds at the choice point (George et al., 2023). All trials with zlnIdPhi scores above 0.3 were examined for verification as VTEs. Using the first visualization method (Figure 2B left), position data from the stem to the choice point exit were plotted with the normalized velocity overlaid. Trials with clear head-sweeping or pausing behavior at the T-intersection were retained as VTEs. Trials with ballistic choice trajectories and/or complex head movements occurring before the choice point entry or after the rat had entered a goal arm were marked as false positive VTE trials. Trials that failed the first inspection were selected for a second round of visualization, when position data were sequentially plotted to “play back” the selected trial. Trials that passed both visualization steps were retained as VTE trials. A second method (Figure 2B right) was used to identify VTE trials with zlnIdPhi scores below 0.3, where high velocity head-sweeping movements could have resulted in a below- threshold zlnIdPhi score and an incorrect classification as a non-VTE trial. This approach determined instances when the rat entered rectangles in both the left and right goal arms of the T-maze during the same trial. These trials were inspected to confirm head-sweeping behaviors at the choice point. Trials that passed this inspection were classified as VTE trials.

**Figure 2.**
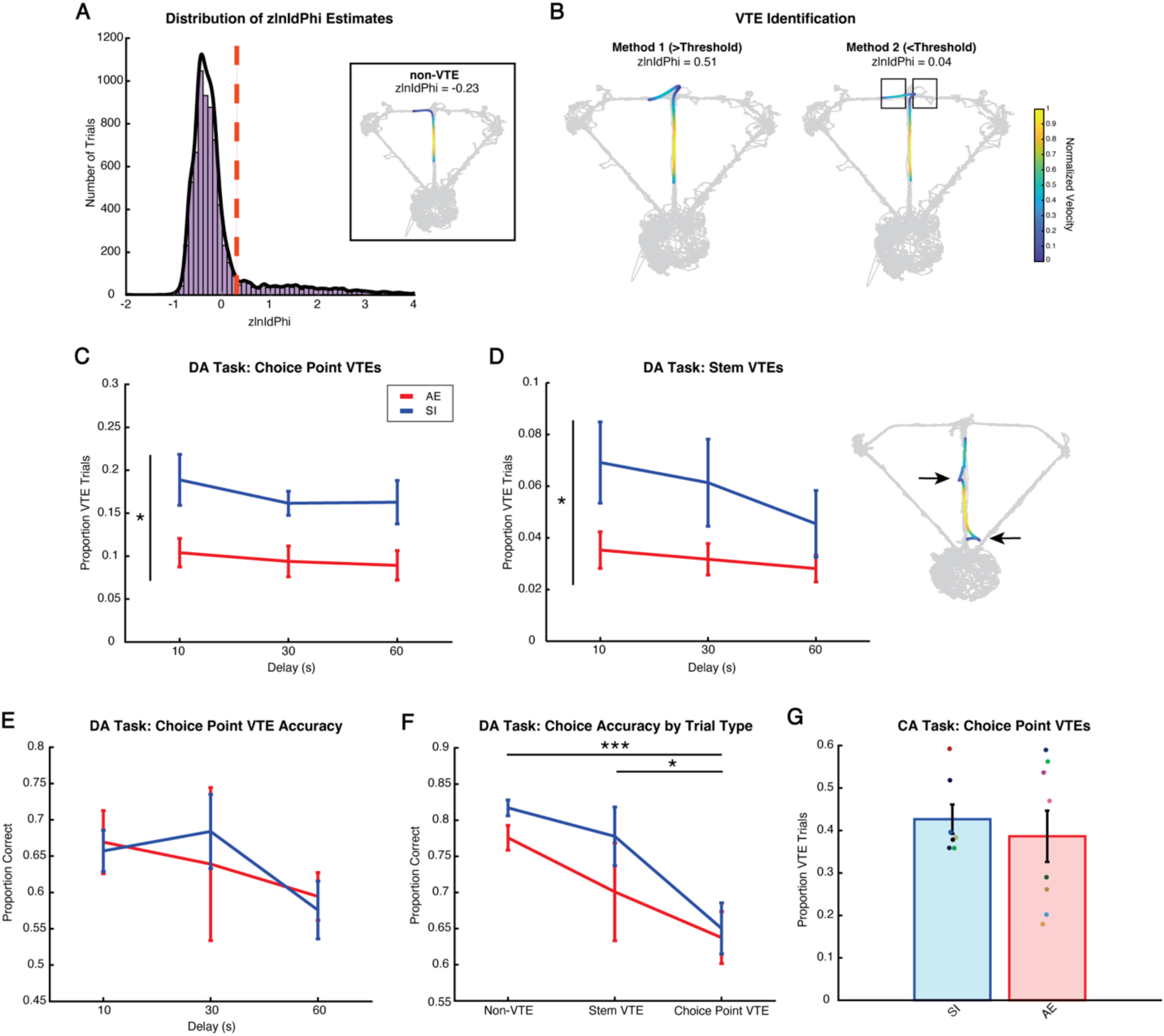
Vicarious trial and error behaviors are less frequent in the alcohol exposed group compared to the sham intubated group. **A)** DA task zlnIdPhi distribution based on choice point tracking data. The VTE threshold (zlnIdPhi= 0.3; red dashed line) was determined as the point where the zlnIdPhi distribution deviated from a normal distribution. Inset: Example non-VTE trial. Trial trajectory overlays tracking data from an example recording session (light gray). Trajectory color represents the normalized velocity of the rat. **B)** Left: Method 1 of VTE trial visualization. Example VTE trial with zlnIdPhi score above threshold. Right: Method 2 of VTE trial visualization. Example VTE trial with zlnIdPhi score below threshold but where the rat enters both goal arms (black boxes). This method allowed us to identify VTE trials that would have originally been excluded due to high velocity through the choice point. Both Method 1 and Method 2 were used to identify VTE trials (see Methods for details). **C)** The overall proportion of trials with a VTE in the choice point is lower in the AE (red) group across delays compared to the SI (blue) group. **D)** Left: The AE group shows fewer VTEs in the stem of the T-maze than the SI group. The proportion of VTE trials is not affected by delay length. Right) Example trial with VTEs (indicated with arrows) in the T-maze stem. **E)** Choice accuracy on the subset of trials with VTEs at the choice point. Compared to choice accuracy on all trials, accuracy on VTE trials did not decrease with increased delay length. **F)** Choice accuracy on non-VTE trials, trials with a VTE in the stem (stem VTE) and trials with a VTE at the choice point (choice point VTE) collapsed across delay. While there was no significant difference in accuracy between the AE and SI groups, there was a main effect of trial type on choice accuracy such that accuracy was significantly lower on choice point VTE trials compared to stem VTE and non-VTE trials. **G)** AE and SI groups show similar proportions of VTE trials at the choice point during the CA task. Colored dots indicate individual rats. *p<0.05, ***p<0.001. Error bars represent mean +/- standard error of the mean.

We also examined VTEs in the T-maze stem. zlnIdPhi of 1.5 was chosen as the threshold value based on the zlnIdPhi distribution generated using stem tracking data from each trial. Trials with above threshold zlnIdPhi scores underwent visualization through Method 1. To examine VTEs at the choice point during the CA task, data underwent both visualization methods, except lnIdPhi scores were not z-scored per rat as there were fewer trials. An lnIdPhi of 4.0 was determined to be the VTE threshold for the CA task.

Analyses examining the proportion of VTEs per session (Figure 4A-B) included data up until session 13, as each recording day contained data from at least half of the rats in each group until this session.

#### DA task choice accuracy across sessions

To examine DA task choice accuracy over testing sessions, 12 implanted rats (7 AE, 5 SI) from the current study were added to an additional dataset consisting of 39 rats (17 AE, 22 SI). These additional rats completed the same experimental procedure as the rats from the current study except that they were not implanted with recording drives. As rats in the previous dataset completed 6 sessions of DA task testing, we analyzed task performance over these sessions in both groups. The sample size accounts for rats excluded from choice accuracy analysis: 5 implanted rats were removed due to recording issues that prevented at least 1 of the first 6 sessions from being completed, 1 implanted rat was determined to be an outlier (greater than 3 scaled median absolute deviations from the median; indicated by a red “X” in Figure 1E; this rat was excluded from all analyses) and 2 rats from the additional dataset were found to have BAC results below 100 mg/dL and were excluded. If the recording headstage became unplugged from implanted rats, the corresponding trial was excluded from the calculation of a choice accuracy score for that session.

#### Perseverative errors

A perseverative error occurred if a rat made an incorrect choice on two consecutive trials of the DA task (ex. left-right-right-right corresponds to correct-error1-error2). The proportion of perseverative errors was calculated as the number of repeated choice errors divided by the total number of errors.

### Electrophysiological analysis

#### Extracting LFPs in the choice point

LFPs were extracted over timestamps when rats occupied the choice point of the T-maze. The 3rd degree polynomial was removed from LFPs using *detrend.m*. The detrended signal was then z-scored to account for overall power distribution differences between rats due to increased signal amplitude after copper mesh cages were introduced to the surgery procedure.

#### Coherence and power spectral density

To examine mPFC and HPC oscillatory activity and the magnitude of mPFC-HPC coupling during choice point occupancy, power spectral density estimates (*pwelch.m)* and magnitude-squared coherence (*mscohere.m)* were calculated over 1-50 Hz at a frequency resolution of 0.5 Hz. Power spectral density is a measure of the power (squared amplitude) of a signal scaled by frequency. The log10 of the power spectral density estimates was taken to account for 1/f noise. Magnitude-squared coherence is a metric that describes the degree to which two signals are temporally correlated and ranges from 0 (no correlation) to 1 (perfect correlation):

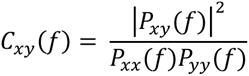

The magnitude-squared coherence (C_xy_) at a specified frequency (f) is the square of the absolute value of the cross-power spectral density (P_xy_) scaled by the power spectral density of each signal (P_xx_, P_yy_). As 1.25 seconds of data is sufficient for reliable estimates of theta coherence (Stout et al., 2023), trials that did not reach this threshold were excluded from the analysis. To account for quick passes through the choice point on non-VTE trials, LFPs were concatenated by session for non-VTE LFP analysis.

A moving window approach was used to examine the prevalence of mPFC-HPC coupling during choice point occupancy on VTE and non-VTE trials. First, LFP signals were concatenated by rat. Magnitude-squared coherence was then calculated from 6-10 Hz at a frequency resolution of 0.5 Hz over 1.25-second time windows (“coherence events”) that were gradually shifted by 250 milliseconds (Stout et al., 2023; Figure 7A). The final samples of each rat’s concatenated signal were excluded as the remaining data samples did not meet the 1.25-second minimum required for inclusion in coherence analysis. The mean scores from each 1.25-second coherence event were compiled into empirical cumulative distribution function (CDF) plots.

### Machine learning analysis

To determine if our data could be used to predict whether a rat belonged to the AE or SI group, we built 2 machine learning algorithms (K-Nearest Neighbors (KNN) Classifier and Euclidean Classifier) using leave-one-out approaches. Features were z-scored to account for scaling differences.

In each iteration using the KNN Classifier, the Euclidean distance of the test data vector (representing one rat) to each vector in the training data (representing every other rat) was calculated and sorted. The 7 nearest vectors (neighbors) were determined (refer to Figure 8A), and the test data was classified as belonging to the group to which at least 4 of 7 of the nearest neighboring rats belonged. To determine if our classifier was performing above chance levels, we tested the classifier 1,000 separate times using shuffled labels of AE and SI rats. A z-test was performed to test if the accuracy distribution generated using the shuffled labels was significantly different from the accuracy score using the actual labels (Sangiamo et al., 2020). In each iteration using the Euclidean Classifier, a vector representing all the data from one rat was removed (test data). The remaining data (training data) were separated by group, and the mean vectors were calculated. The Euclidean distance between the test data and each of the mean vectors was determined, and the test data was then classified as belonging to the group that corresponded to the shortest distance. Accuracy was calculated as the number of correct classifications divided by the total number of iterations (17; each rat was excluded once).

### Statistical analysis

All VTE choice accuracy analyses required a contribution of at least three trials at each level of the independent variable (George et al., 2023). If a rat did not meet this parameter, the rat was excluded from that test. Statistical analysis was conducted in MATLAB or JASP (ANOVAs). Significant ANOVA results (p<0.05) underwent Bonferroni correction for multiple comparisons. Corrected p values will be referred to as p_bonf_. Information regarding statistical tests is stated in each result section. Cohen’s D was calculated with *computeCohen_D.m* by R.G. Bettinardi (MATLAB) or in JASP. Figures were generated in MATLAB and edited in Adobe Illustrator.

### Code Accessibility

Data and code will be made available upon request.

## Results

### Alcohol exposure disrupts choice behaviors

Despite previous reports of impaired executive functioning in our FASD rodent model (Gursky et al., 2021) and impaired spatial working memory in other models of 3rd trimester AE (Thomas et al., 1996, Wozniak et al., 2004), we did not observe a spatial working memory deficit as DA task accuracy did not differ between groups (group: F(1,15)=0.512, p=0.485; delay by group: F(2,30)=0.140, p=0.870; repeated measures ANOVA; N=7 SI rats, 10 AE rats; Figure 1D). The proportion of correct trials decreased with increasing delay in both groups (F(2,30)=42.376, p<0.001, η^2^_p_=0.739; post hoc comparisons: 10-30s t=2.806, p_bonf_=0.026, d=0.758; 10-60s t=8.996, p_bonf_<0.001, d=2.429; 30-60s t=6.190, p_bonf_<0.001, d=1.671; two-sample, two-tailed t-test). While spatial working memory was not disrupted by AE, we found that that the AE group spent significantly less time in the choice point than SI controls (t(15)=2.528, p=0.023, d=1.246, two-sample, two-tailed t-test; N= 7 SI rats, 10 AE rats; Figure 1E). Together, these results indicate that AE altered choice behaviors without disrupting spatial working memory.

### Alcohol exposed rats engage in less vicarious trial and errors than controls on the delayed alternation task

To further characterize how choice behaviors were impacted by AE, we investigated VTEs, which are behaviors associated with flexible decision-making, deliberation, and uncertainty (George et al., 2023; Papale et al., 2012; Redish, 2016; Schmidt et al., 2013). We first examined whether there was a relationship between the proportion of trials with a VTE, working memory demand, and AE (Figure 2C; N=7 SI rats, 10 AE rats). We found a main effect of group on the proportion of trials that had VTEs, with the AE group exhibiting a lower proportion of VTE trials than SI controls (F(1,15)=8.540, p=0.011, η^2^_p_=0.363; repeated measures ANOVA). There was no main effect of delay length or delay by group interaction on the proportion of VTE trials, demonstrating that working memory load did not affect overall VTE occurrence (delay: F(2,30)=2.147, p=0.134; delay by group: F(2,30)=0.319, p=0.729).

While examining tracking data to confirm VTEs at the choice point, we noticed instances of rats displaying VTE-like behaviors on the maze stem (Figure 2D right). We were curious if these “stem VTEs” would also be lower in the AE group compared to the SI group. A repeated measures ANOVA revealed a main effect of group on VTE trial proportion in the stem of the maze, with the AE group showing a lower proportion of trials with stem VTEs than the SI group (F(1,15)=4.583, p=0.049, η^2^_p_=0.234; N=7 SI rats, 10 AE rats; Figure 2D left). There was no effect of delay length or delay by group interaction on VTE proportion in the T-maze stem (delay: F(2,30)=2.995, p=0.065; delay by group: F(2,30)=0.907, p=0.415). Both the SI and AE groups performed a greater proportion of VTEs in the choice point than the stem of the maze (SI: t(6)=8.182, p=0.0002, d=3.093; AE: t(9)=4.581, p=0.001, d=1.449; one-sample, two-tailed t-test against a null of 0; data not shown).

Our findings suggest that developmental AE leads to less deliberation during decision-making in adulthood. However, an alternative explanation is that our results instead reflect a motor impairment (Goodlett et al., 1991; Klintsova et al., 1998; Thomas et al., 1996), as AE during the brain growth spurt also damages the cerebellum (Bonthius & West, 1991; Hamre & West, 1993). To investigate this possibility, we examined VTEs at the choice point during the CA task, a task with a comparatively low working memory demand compared to the DA task. Tracking data were recorded from 8 AE rats and 7 SI rats during 1-5 CA task sessions occurring late in training. In contrast to the DA task, there was no difference in time spent in the choice point or the proportion of trials with VTEs between groups on the CA task (time spent: t(13)=0.062, p=0.951, data not shown; VTE: t(13)=0.556, p=0.587; two-sample, two-tailed t-test; Figure 2G). As the AE group was capable of performing VTEs at similar levels as the SI group, it is unlikely that motor impairments explain VTE differences on the DA task.

We next investigated whether choice accuracy on VTE trials differed between groups and if delay length affected performance on these trials. In contrast to our overall DA task accuracy results, we found that choice accuracy did not change across delays on trials with VTEs at the choice point (F(2,24)=1.531, p=0.237; repeated measures ANOVA; N=7 SI rats, 7 AE rats; Figure 2E). AE and SI groups also performed similarly on choice point VTE trials across delays (group: F(1,12)=0.007, p=0.933; delay by group: F(2,24)=0.236, p=0.792). Due to the low trial count of stem VTEs, we did not analyze the relationship between choice accuracy and delay length.

As VTEs are associated with uncertainty and conflict, they are also related with poorer task performance compared to non-VTE trials (Amemiya & Redish, 2016). We examined whether this relationship was disrupted after AE and how VTE location in the maze (either the stem or the choice point) impacted choice accuracy (Figure 2F). Both non-VTE trials and stem VTE trials showed higher choice accuracy than choice point VTE trials (F(2,26)=10.105, p<0.001, η^2^_p_=0.437; repeated measures ANOVA; post hoc comparisons: non-VTE vs choice point VTE t=4.449, p_bonf_<0.001, d=1.387; stem VTE vs choice point VTE t=2.782, p_bonf_=0.030, d=0.867; two-sample, two-tailed t-test; N= 7 SI rats, 8 AE rats). Interestingly, choice accuracy on stem VTE trials was not significantly different from choice accuracy on non-VTE trials (t=1.667, p_bonf_=0.323; two-sample, two-tailed t-test). Choice accuracy was not affected by AE (group: F(1,13)=1.139, p=0.305; trial type by group: F(2,26)=0.438, p=0.650).

Together, our results suggest that AE during the brain growth spurt leads to reduced deliberative behaviors during HPC-dependent working memory, as the AE group exhibited fewer VTEs on the DA task while groups showed similar amounts of VTEs on the CA task. While VTE frequency was lowered after AE, the AE group did not show a choice impairment on VTE trials. We also found that VTEs were not limited to locations near the T-intersection of the maze, demonstrating that rats occasionally began engaging in these behaviors shortly after trial initiation. Moreover, engaging in deliberation early in the trial (in the stem versus the choice point) may have benefited impending choice accuracy. Choice accuracy on choice point VTEs was not affected by delay, indicating that these behaviors manifested similarly regardless of working memory load.

### Disturbed relationship between vicarious trial and error and choice outcomes following alcohol exposure

VTEs have been shown to be more common on error trials compared to correct trials (Bett et al., 2012; Schmidt et al., 2013; but see Miles et al., 2024). To investigate the relationship between AE, trial accuracy, and delay duration on choice point VTE behaviors, we compared the proportion of VTEs occurring on correct and error trials (Figure 3A) for the 10-, 30-, and 60-second delays in the AE and SI groups (N= 7 SI rats, 10 AE rats). There was no significant 3-way interaction between AE, trial accuracy, and delay (F(2,30)=1.167, p=0.325; repeated measures ANOVA). However, there was a significant interaction between trial accuracy and group, as the proportion of VTE error trials (Figure 3B), but not VTE correct trials (Figure 3C), was lower in the AE group compared to the SI group (trial accuracy by group: F(1,15)=11.316, p=0.004, η^2^_p_=0.430; trial accuracy: F(1,15)=70.124, p<0.001, η^2^_p_=0.824; post hoc comparisons: error AE vs error SI t=-4.730, p_bonf_<0.001, d=1.886; correct AE vs correct SI t=-1.777, p_bonf_=0.537; two-sample, two-tailed t-test). Both groups also performed a greater proportion of VTE error trials than VTE correct trials (correct AE vs error AE t= −3.904, p_bonf_=0.008, d=0.877; correct SI vs error SI t=-7.652, p<0.001, d=2.054; two-sample, two-tailed t-test).

**Figure 3.**
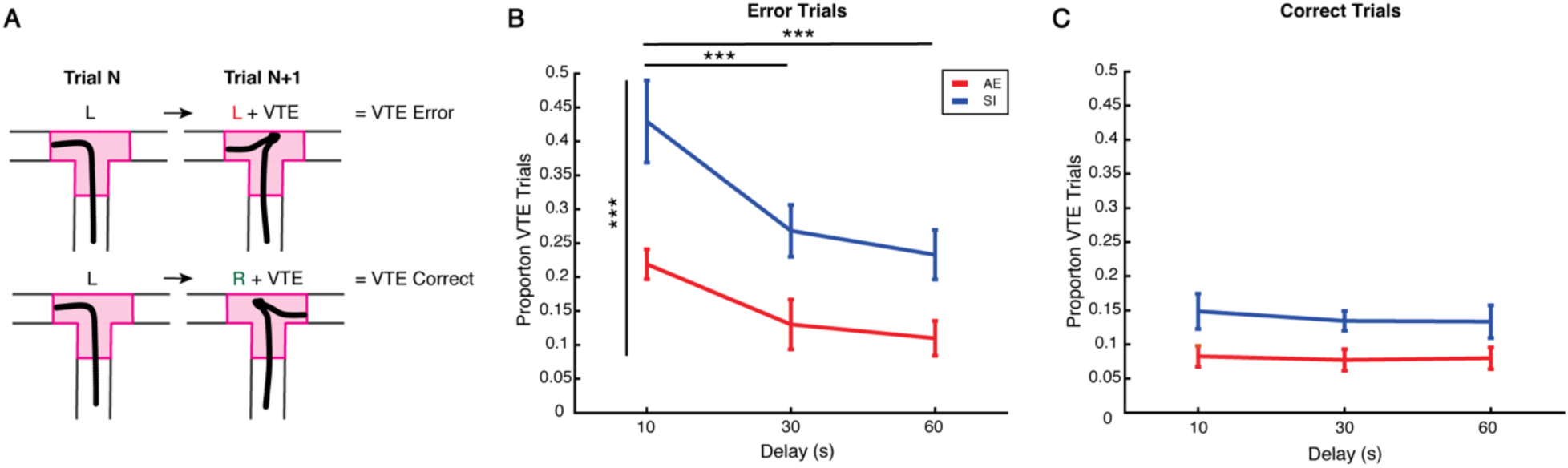
The proportion of error trials with vicarious trial and errors decreases with delay duration and is lower in the alcohol exposed group compared to the sham intubated group. **A)** Schematic of VTE Error (top) and VTE Correct (bottom) trials. Choice point trajectories from example trials are represented in black. L=left choice, R=right choice. The choice point is highlighted in pink. **B)** The proportion of VTE error trials is lower in the AE group (red) compared to the SI group (blue). VTE error trials decrease with delay duration, with the highest proportion of VTEs occurring on 10-second delay error trials. **C)** The proportion of VTE correct trials is not significantly different between groups. VTE trial proportion is not affected by delay on correct trials. **B-C)** VTEs occur on a greater proportion of error trials compared to correct trials. ***p<0.001. Data are represented as mean +/- standard error of the mean.

There was also a significant trial accuracy by delay interaction, with the proportion of VTE error trials decreasing with delay duration and the greatest proportion occurring on 10-second delay trials (trial accuracy by delay: F(2,30)=16.510, p<0.001, η^2^_p_=0.524; delay F(2,30)=14.375, p<0.001, η^2^_p_=0.489; repeated measures ANOVA; post hoc comparisons: 10s error-30s error t=5.977, p_bonf_<0.001, d=1.499; 10s error-60s error t=7.314, p_bonf_<0.001, d=1.834; 30s error-60s error t=1.336, p_bonf_=1.000; two-sample, two-tailed t-test). Therefore, VTE error trials followed an opposite trend to error patterns typically observed in delayed alternation tasks, which increase with delay duration, as reported in our dataset (See Figure 1D) and previous studies that did not separate VTE from non-VTE trials (Ainge et al., 2007; de Mooij-van Malsen et al., 2023; Layfield et al., 2015). In contrast, there was no relationship between the proportion of correct trials with VTEs and delay duration (10s correct-30s correct t=0.464, p_bonf_=1.000; 10s correct-60s correct t=0.428, p_bonf_=1.00, 30s correct-60s correct t=-0.036, p_bonf_=1.000).

Our results indicate that the lower proportion of VTEs exhibited by the AE group (Figure 2C) is likely driven by a reduction in VTEs performed during error trials compared to the SI group. Furthermore, while rats made fewer choice errors on 10-second delay trials compared to 30- and 60-second trials, a higher proportion of these trials had VTEs.

### Altered relationship between experience and vicarious trial and error in the alcohol exposed group

We were next interested in examining if VTE differences between groups were associated with choice accuracy differences at the session level on the DA task. As VTEs are inversely related to learning (Griesbach et al., 1998; Hu & Amsel, 1995; Muenzinger, 1938; Tolman, 1939), we first predicted that the proportion of VTE trials per session would be negatively correlated with choice accuracy. Consistent with previous findings, VTE proportion was negatively correlated with accuracy for both groups (SI: r=-0.3562, p=0.0015; AE: r=-0.3467, p=0.0004; r=correlation coefficient, Pearson’s correlation; N=77 sessions from SI rats, 99 sessions from AE rats; Figure 4A). We also predicted that the greatest proportion of VTEs would occur during the first DA task sessions when rats would need to adjust their strategy to address changes in task demands relative to the CA task and that these behaviors would decrease over sessions. Interestingly, while the SI group demonstrated a reduction in VTE proportion over sessions, this trend was not observed in the AE group, which showed no change in VTE proportion over sessions (SI: r=-0.3806, p=0.0006; AE: r=-0.0569, p=0.576; Pearson’s correlation; N=77 sessions from SI rats, 99 sessions from AE rats; Figure 4B).

**Figure 4.**
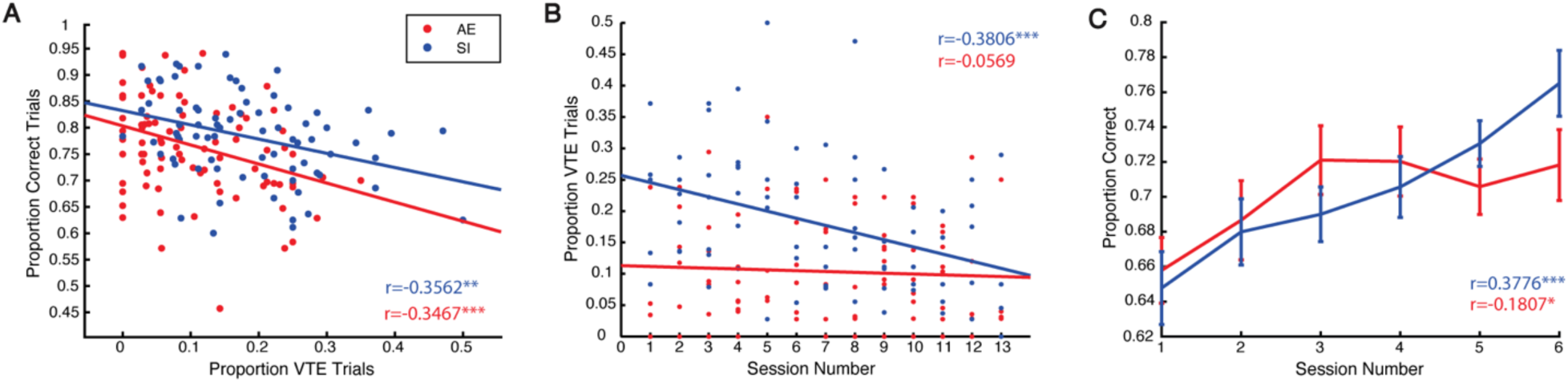
The frequency of vicarious trial and errors decreases with experience in the sham intubated group, but not the alcohol exposed group. **A)** Scatterplot demonstrating a significant negative correlation between the proportion of VTEs and session choice accuracy in the AE (red) and SI (blue) groups. To directly compare the relationship between VTEs and session accuracy, only trials with position data (and therefore VTE data) were included in the calculation of a session choice accuracy average. **B)** While VTE proportion decreases over sessions in the SI group, the AE group does not show a change in VTE frequency. **C)** Choice accuracy is not significantly different between groups across sessions on the DA task. Both the SI and AE groups show improvements across sessions, shown as significant positive correlations between session number and choice accuracy. Data are represented as mean +/- standard error of the mean. *p<0.05, **p<0.01, ***p<0.001.

Given that the proportion of VTE trials was lower in the AE group compared to the SI group and the frequency of VTE trials did not change with experience in the AE group, we predicted that the AE group would show an impairment on the task over sessions. We included rats from a previous dataset that completed the same experimental procedure except DA task testing stopped after session 6 and recording drives were not implanted (combined N= 27 SI rats, 24 AE rats). A repeated measures ANOVA revealed that there was no interaction between group and session and no effect of group on choice accuracy (group by session: F(5,245)=1.664, p=0.144; group: F(1,49)=0.008, p<0.929; Figure 4C). There was a main effect of session on choice accuracy (F(5,245)=7.665, p<0.001, η^2^_p_=0.135). Both SI and AE groups improved across sessions (SI: r=0.3776; p<0.001 AE: r=0.1807, p=0.030; Pearson’s correlation). Together, these results further confirm that although VTE behaviors were disrupted in AE rats, this disruption did not prevent rats from successfully performing and improving on the DA task.

### The functionality of deliberative behaviors is reduced after alcohol exposure

Reorienting behaviors have previously been shown to enhance future decision-making (George et al., 2023). As there was a disturbed relationship between VTEs and performance over sessions in the AE group, we were next interested in determining whether the relationship between VTEs and subsequent performance was also altered. We examined choice accuracy on the trial following a VTE trial and found that the AE group had lower choice accuracy following 10-second delay trials with VTEs compared to the SI group (t(12)=2.508, p=0.028, d=1.295; two-sample, two-tailed t-test; N=7 SI rats, 7 AE rats; Figure 5A). This relationship did not exist when considering non-VTE 10-second delay trials t(15)=0.930, p=0.367; N=7 SI rats, 10 AE rats; Figure 5D). Therefore, the impaired performance of the AE group following 10-second delay trials was not a general characteristic of performance and was specific to trials following VTE trials. In contrast, both groups performed similarly on the trial following 30-second and 60-second delay trials with VTEs (30s: t(12)=-1.016, p=0.330; 60s: t(12)=-1.632, p=0.129; Figure 5B-C). Due to the trial sequence of the DA task, trials following 10-, 30-, and 60-second trials were not evenly distributed (Figure 5E). However, these differences do not explain the impaired performance of the AE group after 10-second VTE trials compared to SI controls, as both groups had similar distributions of delay trials following each type of VTE trial.

**Figure 5.**
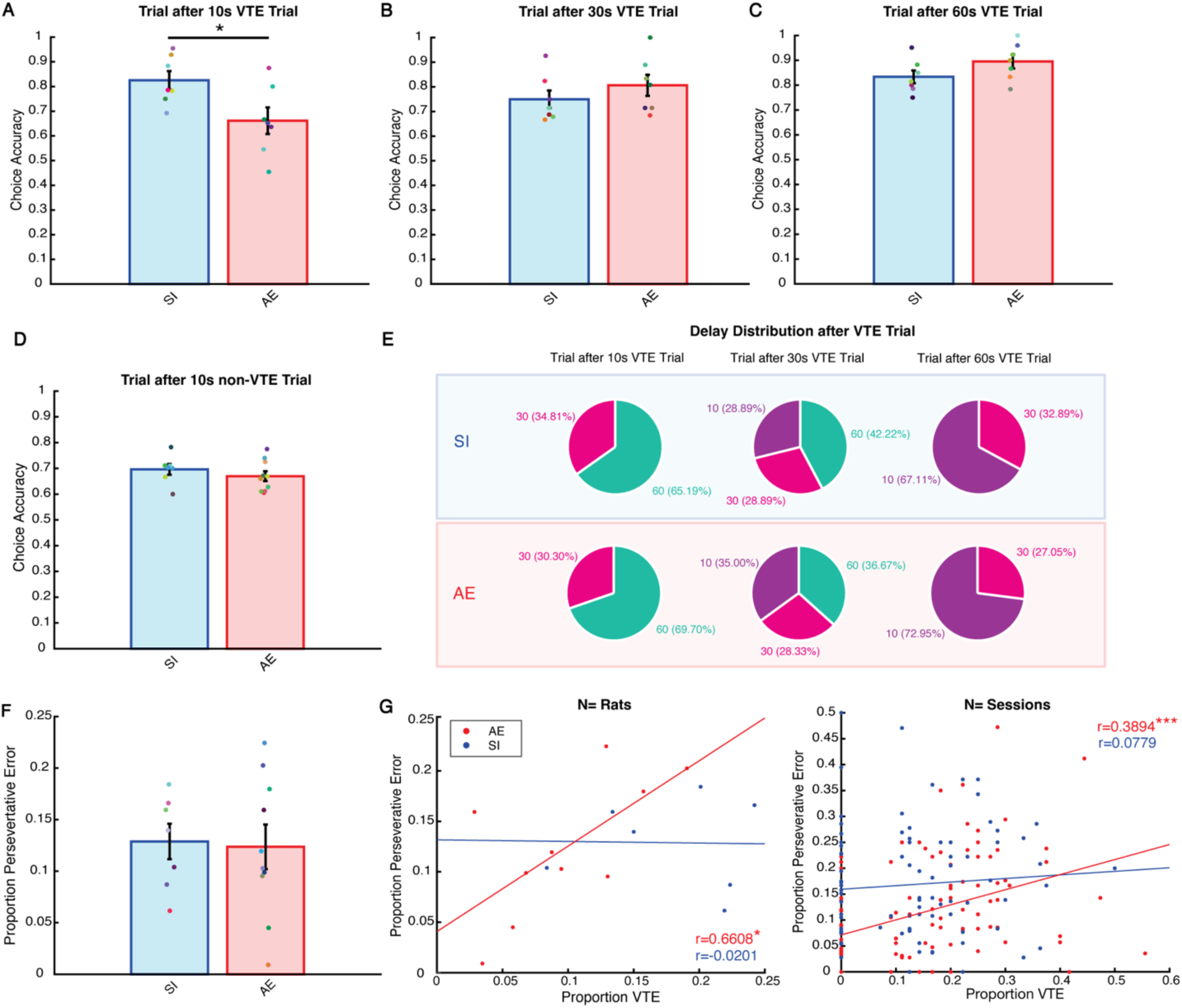
Vicarious trial and errors are associated with perseverative errors in the alcohol exposed group. **A)** The AE group (red) performs poorer on the trial following 10-second delay trials which included VTEs compared to the SI group (blue). In contrast, both groups perform similarly following 30-second **(B)** and 60-second **(C)** delay VTE trials. Colored dots indicate individual rats. **D)** AE and SI groups perform similarly on the trial following 10-second delay trials that were non-VTE trials. **E)** Delay distributions of the trial following a VTE during 10-second, 30-second, and 60-second delay trials. Due to the delay sequence followed during the DA task, 10-second delays and 60-second delays were never followed by consecutive delays of the same length. Delay distributions of the SI group are boxed in blue and delay distributions of the AE group are boxed in red. 10-second delay trials are indicated in purple, 30-second delay trials are indicated in pink, and 60-second delay trials are indicated in green. **F)** Rats in the SI and AE groups perform similar proportions of perseverative error trials during the DA task. **G)** The proportions of VTEs and perseverative errors are positively correlated at the rat (left) and session (right) levels in the AE group (red) but not the SI group (blue). **\***p<0.05. ***p<0.001. Bar plots represent the mean +/- standard error of the mean.

As these results indicated that flexibility may be impaired in AE rats, we decided to investigate measures of executive dysfunction. Inactivation of the mPFC (G.-W. Wang & Cai, 2006), Re (Stout et al., 2022; Viena et al., 2018), and HPC (Hallock et al., 2013) is associated with choice inflexibility, reflected as an increase in repeated choice errors, known as perseverative errors. Similarly, rodent models of third trimester AE have shown increased perseverative errors during spatial working memory and serial spatial discrimination reversal tasks (Thomas et al., 1996, 1997). These findings posed the possibility that we may see an increase in inflexible choice behaviors in our 3rd trimester FASD rodent model. However, we found that the AE and SI groups engaged in a similar proportion of perseverative errors during the DA task (t(15)=0.175, p=0.864; two-sample, two-tailed t-test; N= 7 SI rats, 10 AE rats; Figure 5F).

Given previous work has shown that nucleus reuniens inactivation increases VTEs during perseverative error sequences (Stout et al., 2022), we decided to investigate the relationship between the proportion of perseverative errors and the proportion of VTEs from each rat’s recording sessions. Interestingly, perseverative errors were positively correlated with VTEs in the AE group only, suggesting that AE altered performance such that flexible decision-making behaviors became associated with inflexible decision-making behaviors (individual rats: SI (7 rats) r=-0.0201, p=0.9659; AE (10 rats) r=0.6608, p=0.0375; individual sessions: SI (90 sessions) r=0.0779, p=0.4654; AE (121 sessions) r=0.3894, p<0.001; Pearson’s correlation; Figure 5G). Collectively, these findings suggest that VTE efficacy has been reduced in AE rats as they did not facilitate a flexible choice strategy as reflected in SI controls.

### Alcohol exposure alters mPFC theta oscillations and HPC beta oscillations

mPFC-HPC theta synchrony via the nucleus reuniens has been implicated in decision-making (Hallock et al., 2016) and VTE behaviors (Stout et al., 2022). Therefore, we were interested in examining the effects of AE on mPFC and HPC physiology and synchrony in the theta band (6-10 Hz) during VTEs (Figure 6A). 7 AE and 4 SI rats were included in LFP analysis after verifying electrode placements. The power spectral densities of mPFC and HPC LFPs recorded during choice point occupancy in both the AE and SI groups are shown as a function of frequency in Figure 6B-C (left). We found that theta power in the mPFC was significantly lower in the AE group compared to the SI group during VTEs (t(9)=2.534, p=0.032, d=1.588; two-sample, two-tailed t-test; Figure 6B middle). To determine if this effect was specific to VTEs, we next examined non-VTE trials. After outlier removal, we found that mPFC theta power was also significantly lower in the AE group compared to the SI group during non-VTE trials (t(7)=2.716, p=0.030; d=1.822; N=4 SI rats, 5 AE rats; Figure 6E left). Follow-up analysis revealed that the proportion of VTE trials performed by rats in the AE group, but not the SI group, was negatively correlated with mPFC theta power during VTE trials, but not non-VTE trials (VTE: AE r=0.7843, p=0.0368; SI r=0.4094, p=0.5906; Pearson’s correlation; Figure 6H; non-VTE: AE r=-0.2895, p=0.6366; SI: r=0.1588, p=0.8412; data not shown). In contrast, HPC theta power and mPFC-HPC theta coherence were not different between groups during VTEs and non-VTEs (VTE power: t(9)=0.291, p=0.778; Figure 6C middle; VTE coherence: t(9)=-0.232, p=0.822; Figure 6D middle; non-VTE power: t(9)=0.930, p=0.376; Figure 6F left; non-VTE coherence: t(9)=0.227, p=0.826; Figure 6G left).

**Figure 6.**
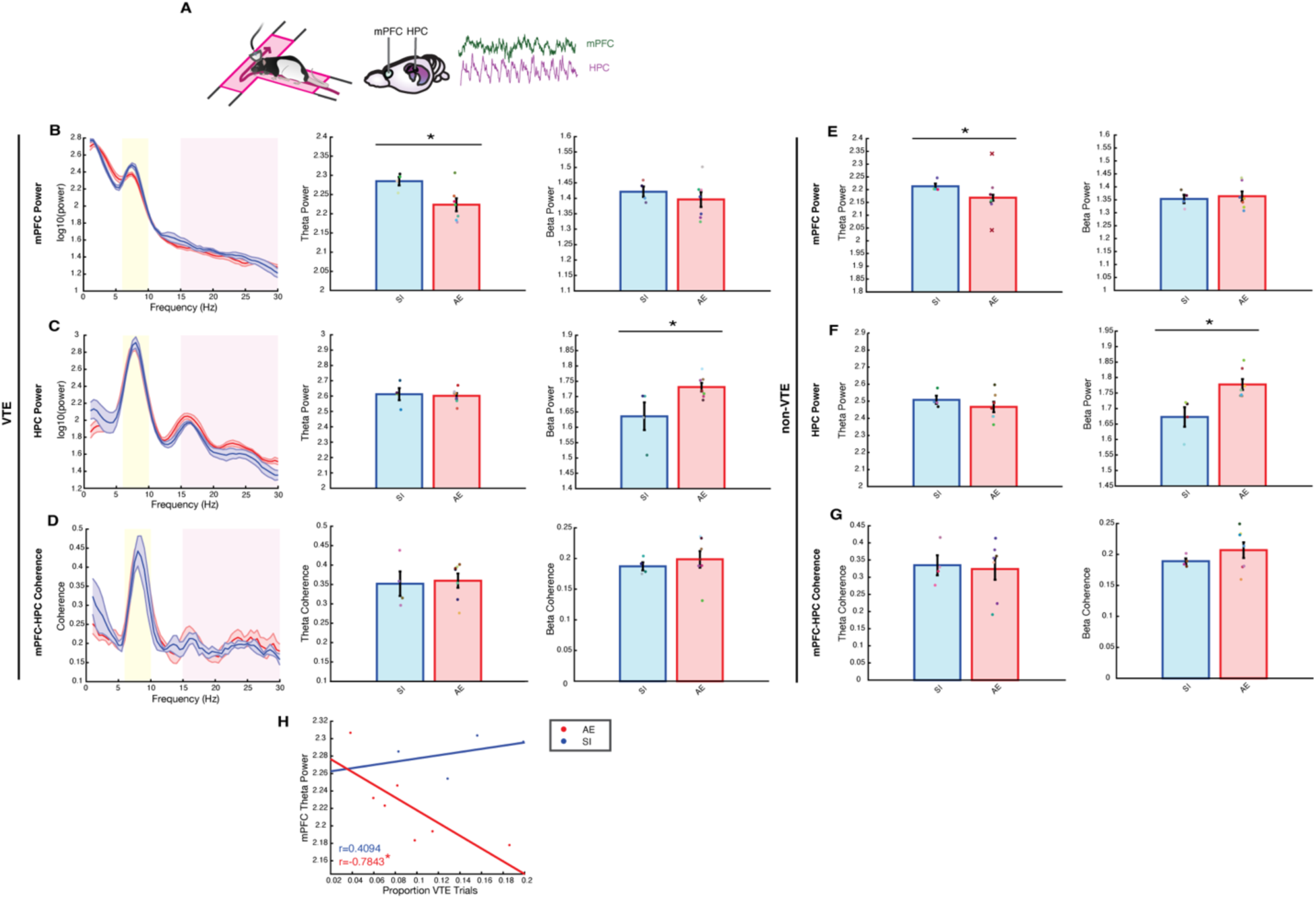
mPFC theta power and HPC beta power are altered after alcohol exposure. **A)** LFPs were recorded from the mPFC and HPC during choice point occupancy (highlighted in pink) on VTE trials. Example signals from the mPFC (green) and HPC (purple) are shown to the right. **B)** Left: mPFC power distribution as a function of frequency for the AE (red) and SI (blue) groups. The mean power distribution is represented as a solid line and the standard error of the mean is represented as the shaded area around the mean. Analyses were performed over the 6-10 Hz theta range (highlighted in yellow) and the 15-30 Hz beta range (highlighted in pink). Middle: Bar plot demonstrating mPFC theta power during VTEs is lower in the AE group compared to the SI group. Right: Bar plot showing mPFC beta power during VTEs is not different between groups. Bar plots represent the mean +/- standard error of the mean. Colored dots indicate individual rats. **C)** Same as B, except for HPC power. HPC theta power is not different between groups (middle), while beta power is higher in the AE group compared to the SI group (right). **D)** Same as B, except for mPFC-HPC coherence. mPFC-HPC theta (middle) and beta (right) coherence are not different between groups. **E)** Left: mPFC theta power is lower in the AE group compared to the SI group during non-VTE trials. Two outlier rats were identified in the AE group and are indicated with a red “X”. Right: mPFC beta power is not different between groups during non-VTE trials. **F)** HPC theta power is not significantly different between groups during non-VTE trials (left), whereas HPC beta power is higher in the AE group than the SI group (right). **G)** mPFC-HPC theta (left) and beta (right) coherence are not different between groups during non-VTE trials. **H)** Scatterplot showing that the proportion of VTE trials is negatively correlated with mPFC theta power during VTE trials in the AE group but not the SI group. Only trials with clean LFP data were considered in the calculation of VTE trial proportion. *p<0.05.

Beta rhythms (15-30 Hz) have also been associated with VTEs (Miles et al., 2024) and synchronize in the mPFC-nucleus reuniens-HPC circuit during memory tasks (de Mooij-van Malsen et al., 2023; Jayachandran et al., 2022). We found that HPC beta power was significantly higher in the AE group compared to the SI group during both VTE and non-VTE trials (VTE: t(9)=-2.520, p=0.033, d=1.580; Figure 6C right; non-VTE: t(9)=-3.188, p=0.011; d=1.998; Figure 6F right). Conversely, mPFC beta power and mPFC-HPC beta coherence during VTEs and non-VTEs were not significantly different between groups (VTE power: t(9)=0.731, p=0.483, Figure 6B right; VTE coherence: t(9)=-0.497, p=0.631; Figure 6D right; non-VTE power: t(9)=-0.372, p=0.719; Figure 6E right; non-VTE coherence: t(9)=-1.024, p=0.333; Figure 6G right).

Together, these results suggest that AE during the brain growth spurt alters mPFC theta rhythms and HPC beta rhythms during both VTEs and non-VTEs without disrupting the magnitude of mPFC-HPC synchrony.

### mPFC-HPC theta coupling events are less common after alcohol exposure

Our results suggested that the magnitude of mPFC-HPC theta synchrony during decision-making was not different between groups. It remained possible that AE could disturb the commonality of mPFC-HPC coupling events, rather than the magnitude. For example, magnitude coherence measures over choice point occupancy could have masked differences in how frequently the mPFC and HPC synchronized over shorter timescales. Therefore, we next used a moving window approach to calculate mPFC-HPC theta coherence over 1.25-second “coherence events” (refer to Methods; example trials with similar magnitude coherence and different coherence event distributions are shown in Figure 7A). We first validated this approach by replicating our previous magnitude coherence results from Figure 6D using the mean coherence magnitude across events for each rat (t(9)=1.129, p=0.288; two-sample, two-tailed t- test; N= 4 SI rats, 7 AE rats; Figure 7B).

**Figure 7.**
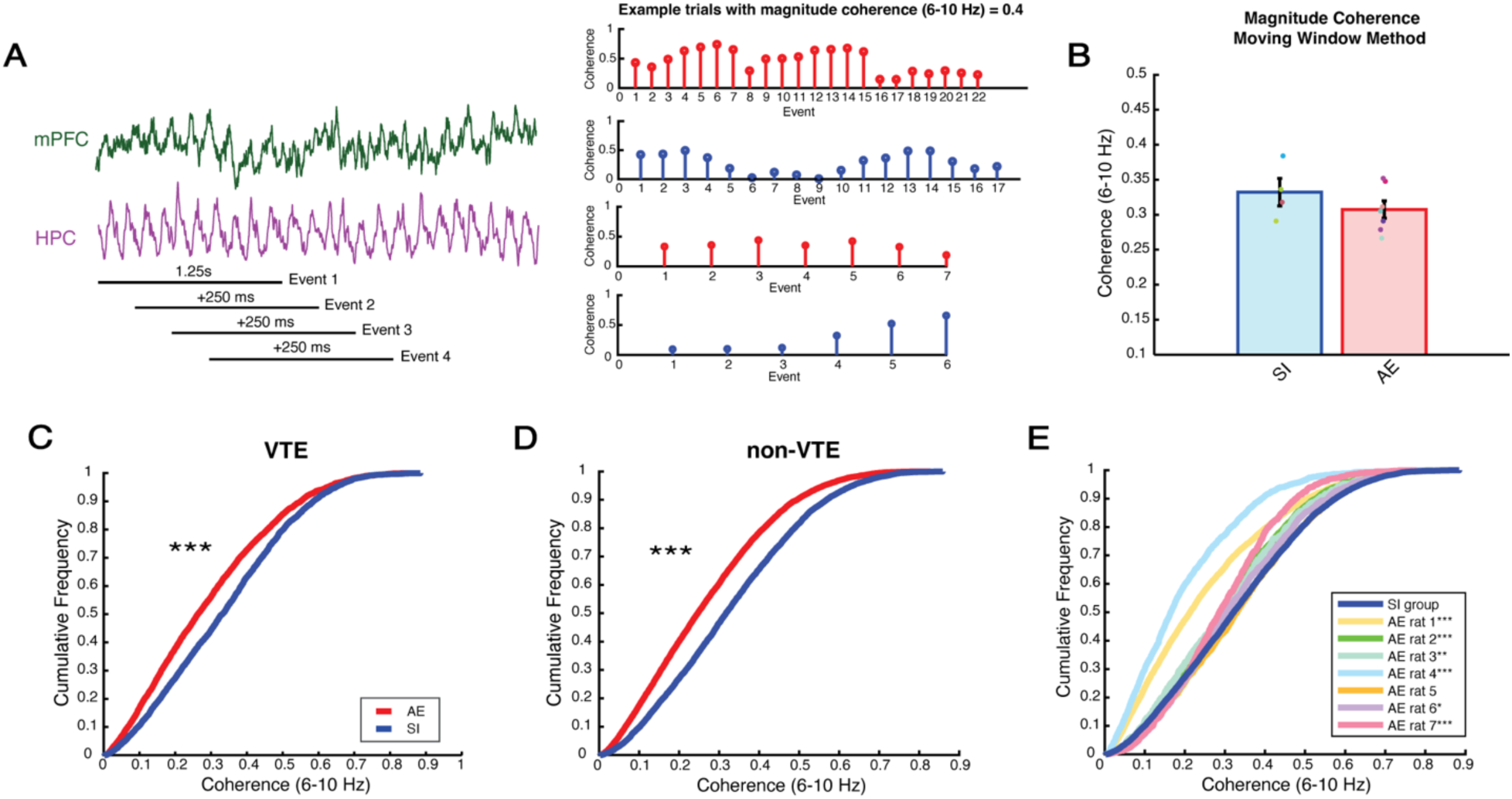
The prevalence of mPFC-HPC synchronous events is altered after alcohol exposure. **A)** Left: Schematic of moving window method to calculate mPFC-HPC coherence. Coherence was calculated over 1.25-second (s) events that were gradually shifted by 250 milliseconds (ms). Example LFPs from the mPFC and HPC are represented in green and purple, respectively. Right: Stem plots showing theta coherence across events from example trials of different AE (red) and SI (blue) rats. Each trial has a magnitude coherence of 0.4 during choice point occupancy. Note that the degree of mPFC- HPC synchronization varies across events within this period. **B)** Bar plot demonstrating that magnitude coherence (6-10 Hz) is not different between the AE (red) and SI (blue) groups when calculated with the moving window approach. Colored dots indicate individual rats. **C)** CDF plot showing that the distributions of mPFC-HPC theta coherence events (6-10 Hz) are significantly different between the AE (red) and SI (blue) groups during VTEs. **D)** Same as C, except coherence events were measured from non-VTE trials. **E)** CDF plot showing theta coherence event distributions of individual AE rats compared to the theta coherence event distribution of the SI group (dark blue line). Asterisks in the legend indicate that the coherence event distribution of the corresponding rat is significantly different from the coherence event distribution of the SI group. *p<0.05, **p<0.01, ***p<0.001.

Interestingly, whereas magnitude coherence was not different between groups using either approach, the distributions of theta coherence events were significantly different between groups (k=0.124, p<0.001; two-sample Kolmogorov-Smirnov Test; Figure 7C). The coherence event distribution of the AE group was shifted leftward compared to the SI group, suggesting that AE led to less frequent mPFC-HPC theta coupling. This effect was not specific to VTE trials, as we also observed differences between groups in the distributions of theta coherence events during non-VTE trials (k=0.148, p<0.001; Figure 7D). We noticed variability in the number of coherence events that each AE rat contributed to the overall coherence event distribution. To confirm that these effects could also be observed at the rat level, we then collapsed theta coherence events across VTE and non-VTE trials for each rat and tested the distributions of each AE rat against the SI group distribution. We found that 6/7 AE rats showed coherence event distributions that were significantly different from the SI group distribution (rat 1: k=0.216; p<0.001; rat 2: k=0.083; p<.001; rat 3: k=0.079; p=0.0013; rat 4: k=0.335; p<0.001; rat 5: k=0.016; p=0.462; rat 6: k=0.046; p=0.025; rat 7: k=0.141; p<0.001; Figure 7E). Furthermore, the majority of AE rats demonstrated leftward-shifted distributions, indicating that mPFC-HPC theta coupling events were less common compared to the SI group. Collectively, these results indicate that the incidence of mPFC-HPC synchronous events, but not the magnitude of synchrony, is altered after AE and that this alteration is not specific to VTE trials.

### Using machine learning to predict treatment of alcohol exposed and sham intubated rats

We were next interested in determining whether we could predict the treatment of each rat (as AE or SI) using machine learning. Features consisted of the following categories: time spent in choice point, proportion of VTE trials in the choice point by delay, proportion of VTE trials in the stem by delay, proportion of VTE error trials in the choice point by delay, proportion of VTE correct trials in the choice point by delay, proportion of perseverative error trials, and proportion correct by delay (all results included a full sample size for analysis of 10 AE rats and 7 SI rats, therefore LFP data was excluded). We built a KNN classifier with K=7, as this value resulted in the highest accuracy while considering the fewest neighbors (Figure 8A). We found that postnatal treatment as AE or SI was predicted with above-chance accuracy, correctly classifying 9/10 AE rats and 4/7 SI rats (overall accuracy 76.5%; z=2.393, p=0.017; one-sample, two-tailed z-test; Figure 8B). To further validate our KNN Classifier, we also built a Euclidean Classifier to predict treatment. Similar to the KNN Classifier, the Euclidean Classifier correctly predicted 9/10 AE rats and 4/7 SI rats, performing at an identical accuracy of 76.5% (Figure 8C). These results demonstrate our classifiers could reliably predict whether a rat was exposed to alcohol during development based on behavioral data from the DA task.

**Figure 8.**
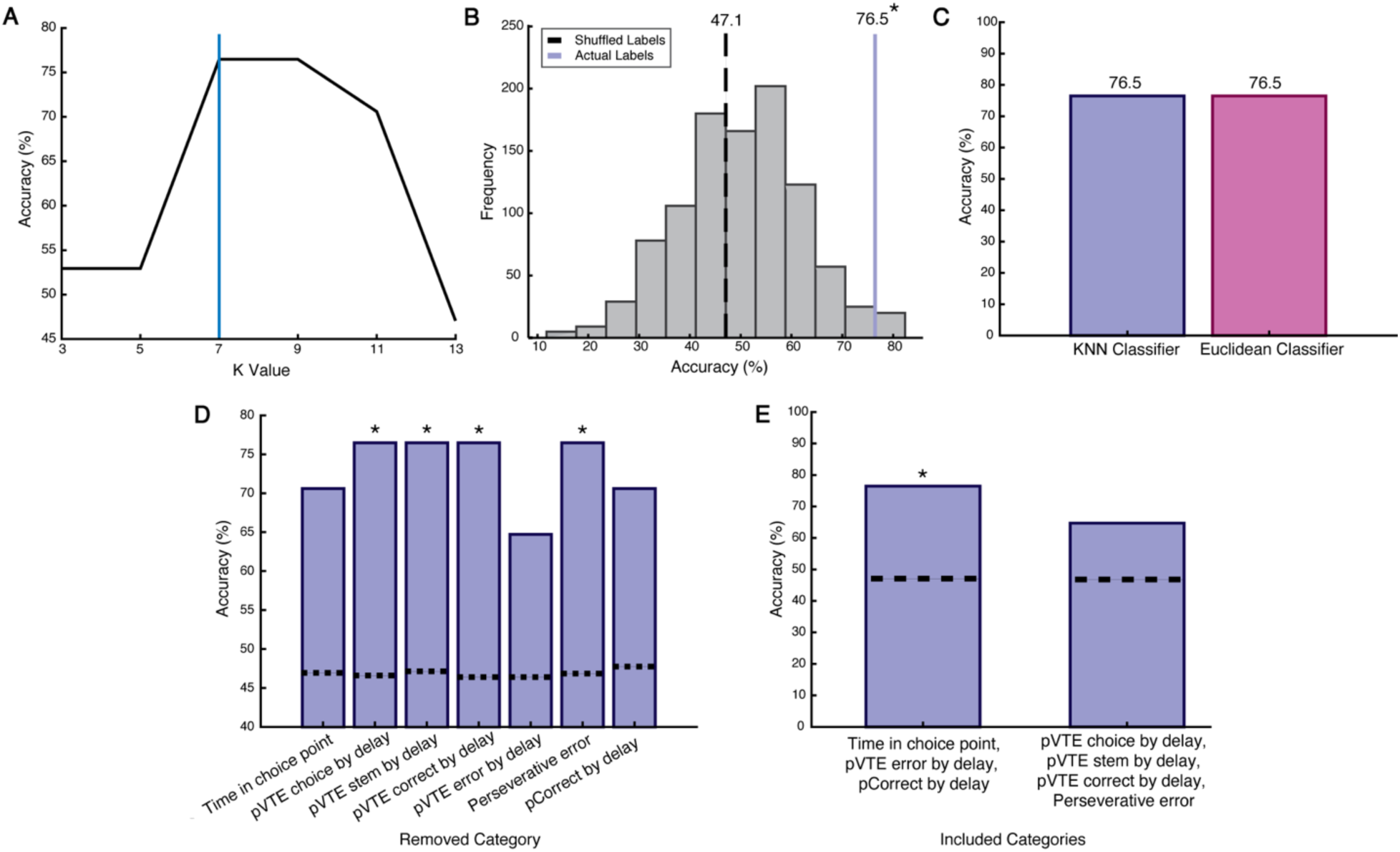
Using supervised machine learning to predict treatment with behavioral data. **A)** Determining K for KNN Classifier. As accuracy initially plateaus at 7 (blue line), this value was chosen as K. **B)** The KNN Classifier performs at 76.5% accuracy (purple line) when using the true labels to predict the treatment of AE and SI rats. Accuracy is significantly above chance levels as determined by performing a z-test with the accuracy distribution obtained by shuffling the labels of AE and SI rats (mean accuracy represented with a black dashed line). **C)** Both a KNN Classifier (purple) and a Euclidean Classifier (pink) achieve the same accuracy when predicting the treatment of AE and SI rats. **D)** Iteratively removing categories from the KNN Classifier to determine which categories contribute to accurate classification. Time spent in the choice point, the proportion of VTE error trials by delay, and overall choice accuracy on the DA task by delay are the only categories that when removed result in reduced classifier accuracy (not significantly different from chance levels). Dashed lines represent mean accuracy of the shuffled distributions. **E)** Using only time spent in the choice point, the proportion of VTE error trials, and overall choice accuracy, the classifier performs at identical accuracy as when all categories are included (C) and performs significantly above chance levels. Testing the classifier on all remaining categories results in accuracy that is not significantly different from chance levels. pVTE=proportion of VTE trials. pCorrect= proportion of correct trials. *p<0.05 compared to shuffled distribution.

To determine which combination of behaviors could characterize a potential phenotype for third trimester AE in our model, we identified which categories were most important for the correct classification of rats as AE or SI. We then iteratively removed each category from the KNN classifier, determined accuracy, and found that classifier accuracy decreased below chance levels (p>0.05) only when time spent in the choice point, the proportion of VTE error trials at each delay, or proportion correct at each delay were excluded (Figure 8D). Using solely these three categories to predict treatment, the classifier again performed at an accuracy of 76.5%, which was above chance levels (z=2.258, p=0.024; Figure 8E). In contrast, when all remaining categories were used to predict treatment, the classifier performed below chance levels (z=1.345, p=0.179).

These results demonstrate that only three categories were required for our classifier to reach peak accuracy. In addition, while accuracy on the DA task was not significantly different between groups (Figure 1D), its interaction with time spent in the choice point (Figure 1E) and the proportion of VTE error trials (Figure 3B) was important in characterizing a phenotype of AE versus SI rats.

## Discussion

In this study, we show that AE during the brain growth spurt led to disrupted choice behaviors and altered mPFC-HPC physiology and connectivity without impairing spatial working memory, suggesting a selective disruption to executive function following AE. We further demonstrate that a machine learning algorithm could predict whether rats were AE based on behavioral measures from our task, identifying a phenotype for our model of third trimester AE.

The AE group performed a lower proportion of VTEs in the choice point and stem of the T-maze compared to the SI group during the DA task. There was no difference in VTE proportion between groups on the CA task, showing that AE rats can perform VTEs normally during a task that has a low working memory demand and does not require HPC (Ainge et al. 2007). In contrast, the DA task has a working memory component that increases with delay duration, is HPC-dependent (Ainge et al. 2007), and relies on mPFC-HPC interactions via the nucleus reuniens (Hallock et al., 2016), particularly during VTEs (Stout et al., 2022). Consequently, the lower proportion of VTEs in the AE group compared to the SI group is likely due to AE-related dysfunction within this circuit disrupting processes underlying deliberation.

Although VTEs were reduced, spatial working memory was unimpaired in the AE group. This result was surprising given reductions in VTEs are associated with learning and memory deficits (Bett et al., 2012; Blumenthal et al., 2011; Griesbach et al., 1998; Hu & Amsel, 1995; Kidder et al., 2021). VTEs were also unrelated to task acquisition in the AE group, indicating that the AE group utilized a strategy that relied less on VTEs but was effective in making correct choices. In contrast, in agreement with previous studies, the SI group showed a reduction in VTEs across sessions that coincided with increased choice accuracy, suggesting rats utilized a deliberative strategy upon task introduction that became less necessary as proficiency increased (Griesbach et al., 1998; Hu & Amsel, 1995; Muenzinger, 1938; Redish, 2016; Tolman, 1939). To our knowledge, we are the first to show that VTE frequency and accuracy are unaffected by working memory demand, which likely relates to VTEs reflecting uncertainty (Amemiya & Redish, 2016; Schmidt et al., 2013). We also demonstrate that VTEs in the T-maze stem had higher accuracy than VTEs in the choice point, indicating that the timing of VTEs relative to the choice has implications for subsequent decision-making.

The AE group performed fewer VTE error trials than the SI group, yet both groups committed a similar proportion of choice errors. These results reveal a fundamental difference in choice behavior during error trials following AE. This prediction is supported by our finding that removing VTE error trials as a category from our KNN Classifier resulted in the greatest decrease in classifier accuracy. As VTEs are thought to reflect the evaluation of choices during indecision (Redish, 2016), our results suggest that the AE group was less likely to engage in deliberative behaviors when uncertain. However, while VTEs were disrupted in the AE group, these behaviors appeared to reflect similar processes in both groups. For example, VTEs were associated with error trials (Bett et al., 2012; Schmidt et al., 2013) and had lower accuracy than non-VTE trials (Amemiya & Redish, 2016). Sessions with high proportions of VTEs also tended to have low choice accuracy (Griesbach et al., 1998; Hu & Amsel, 1995; Tolman, 1939).

Working memory and VTEs were related during choice errors. While a shorter inter-trial delay results in an easier version of the task, it also increases the potential for interference between trials. A recent study found that reorienting behaviors similar to VTEs increased on the delayed non-match to place task when rats were blocked from alternating on the sample phase of the current trial relative to the choice phase of the preceding trial, even though the alternation rule was irrelevant during the sample phase (George et al., 2023). In the current context, rats may have been unable to dissociate the previous from the current trial after 10-second delays, and this conflict could have resulted in VTE occurrence and choice error. As proactive interference decreases with increased inter-trial delay (Grant, 1981), interference would have been less likely after 30- and 60-second delays. Collectively, errors after 10- second delays may be more reflective of interference rather than forgetfulness, whereas the latter may play a larger role in errors after 30- and 60-second delays. We propose that VTEs after the 10-second delay emerge in part due to this interference, whereas VTEs following 30- and 60-second delays arise due to uncertainty. This prediction may explain the choice deficit following 10-second delay VTE trials, but not 30- or 60-second delay VTE trials in the AE group, as VTEs performed to resolve interference rather than deliberate choice options may have separate implications for upcoming behavior.

We also observed that perseverative errors and VTEs were positively correlated in the AE group. The relationship between flexible (VTE) and inflexible (perseverative error) choice behaviors is contradictory but agrees with previous findings of increased VTEs during perseverative error sequences after nucleus reuniens inactivation (Stout et al., 2022). As the AE group did not demonstrate a greater proportion of perseverative error sequences compared to controls, this altered relationship relates to a reduction in the effectiveness of VTE behaviors rather than an increase in inflexible behaviors. mPFC- nucleus reuniens-HPC circuit dysfunction likely contributes to the dissociation between VTEs and flexible decision-making in the AE group given that disrupting these regions affects both VTE and perseverative error behaviors (G.-W. Wang & Cai, 2006; Hallock et al., 2013; Hu & Amsel, 1995; Kidder et al., 2021; Stout et al., 2022; Viena et al., 2018).

In support of circuit disruption following AE, we found that mPFC theta rhythms and HPC beta rhythms were altered during both VTE and non-VTE trials in the AE group compared to the SI group. Theta and beta rhythms have been implicated in VTEs, as both are increased in the mPFC during VTE trials compared to non-VTE trials (Miles et al., 2024) and theta is present in the HPC during VTEs (Amemiya & Redish, 2016; Johnson & Redish, 2007). As disrupting the circuitry involved in VTEs has been shown to alter mPFC and HPC physiology and affect VTE behavior (Schmidt et al., 2019; Stout et al., 2022), changed oscillatory activity in the theta and beta ranges may contribute to altered VTE functionality in our FASD model. This prediction is supported by our finding that mPFC theta power was negatively correlated to the proportion of VTEs performed by rats in the AE group, linking mPFC dysfunction to the observed VTE deficit. These neurophysiology results may further relate to executive functioning deficits previously described in our rodent model (Gursky et al., 2021).

The prevalence of mPFC-HPC synchronous events was also altered in the AE group compared to the SI group. Interestingly, whereas the magnitude of mPFC-HPC theta coherence was not different between groups, our results indicate that mPFC-HPC theta coupling events were less common in the AE group compared to the SI group. Altered mPFC-HPC theta coupling was not specific to VTE trials, suggesting that these changes are a characteristic of mPFC-HPC functional connectivity after developmental AE. It is possible that reorganization within the brain after AE conserved the magnitude of mPFC-HPC synchrony, but not the incidence of synchronous events. We suspect these changes may have conserved spatial working memory but disrupted aspects of decision-making, such as VTEs becoming less effective for AE rats.

In further demonstration that AE during the brain growth spurt has robust effects on behavior in adulthood, our KNN classifier was effective in identifying whether rats were AE using behavioral measures from task performance. We found that time spent in the choice point, VTE error trials, and the proportion of correct trials were most important in accurately classifying rats as belonging to the AE or SI group, and therefore may be among the measures that characterize the FASD phenotype after third trimester AE.

Our results suggest that the AE group occasionally attempted to deliberate. However, disruptions to mPFC-HPC circuitry may have impaired the ability to engage in VTEs when rats performed a task that relied on the integrity of this circuit. Moreover, as the AE group was sometimes unable to utilize these behaviors to inform future decision-making, the benefit of VTEs as a flexible choice strategy was reduced, which could have diminished the need to perform these behaviors as frequently as controls. These factors could have promoted a strategy that did not require VTEs and spared working memory in the AE group.

Collectively, these findings contribute to a better understanding of the effects of third trimester AE on decision-making by providing evidence for behavioral disruptions and neurophysiological alterations within the mPFC-HPC circuit that offer insight into executive functioning deficits after prenatal AE. These results further identify the mPFC-HPC network as a target for therapeutic interventions in FASD patients.

## Author contributions

H.L.R. collected data, performed analysis, and wrote the manuscript. S.K. generated rats and collected data. J.J.S. and A.L.G. provided analysis feedback. A.K. guided animal generation. A.K. and A.L.G. acquired funding. All authors conceptualized questions and contributed to the writing of this manuscript.

## Acknowledgments

This study was funded by the National Institute on Alcohol Abuse and Alcoholism (R01AA027269). We thank Z. Gemzik, K. Matiz, A. Sonchen, and S. Weinstein for technical assistance, I. Smith for assistance with animal generation, and J. Schwarz for analysis advice. We would also like to thank the Office of Laboratory Animal Medicine for their help. The rat cartoon in Figure 1 was created by S. Park. The rat and brain cartoons in Figure 6 were created by G. Costa and W. Tang, respectively, and were downloaded from SciDraw.io.

## References

Ainge, J. A., van der Meer, M. A. A., Langston, R. F., & Wood, E. R. (2007). Exploring the role of context-dependent hippocampal activity in spatial alternation behavior. Hippocampus, 17(10), 988–1002. 10.1002/hipo.20301

Amemiya, S., & Redish, A. D. (2016). Manipulating Decisiveness in Decision Making: Effects of Clonidine on Hippocampal Search Strategies. The Journal of Neuroscience, 36(3), 814. 10.1523/JNEUROSCI.2595-15.2016

Benchenane, K., Peyrache, A., Khamassi, M., Tierney, P. L., Gioanni, Y., Battaglia, F. P., & Wiener, S. I. (2010). Coherent Theta Oscillations and Reorganization of Spike Timing in the Hippocampal-Prefrontal Network upon Learning. Neuron, 66(6), 921–936. 10.1016/j.neuron.2010.05.013

Bett, D., Allison, E., Murdoch, L., Kaefer, K., Wood, E., & Dudchenko, P. (2012). The neural substrates of deliberative decision making: Contrasting effects of hippocampus lesions on performance and vicarious trial-and-error behavior in a spatial memory task and a visual discrimination task. Frontiers in Behavioral Neuroscience, 6. https://www.frontiersin.org/articles/10.3389/fnbeh.2012.00070

Blumenthal, A., Steiner, A., Seeland, K., & David Redish, A. (2011). Effects of pharmacological manipulations of NMDA-receptors on deliberation in the Multiple-T task. Neurobiology of Learning and Memory, 95(3), 376–384. 10.1016/j.nlm.2011.01.011

Bonthius, D. J., & West, J. R. (1991). Permanent neuronal deficits in rats exposed to alcohol during the brain growth spurt. Teratology, 44(2), 147–163. 10.1002/tera.1420440203

Churchwell, J. C., & Kesner, R. P. (2011). Hippocampal-prefrontal dynamics in spatial working memory: Interactions and independent parallel processing. Behavioural Brain Research, 225(2), 389–395. 10.1016/j.bbr.2011.07.045

Coles, C. (1994). Critical Periods for Prenatal Alcohol Exposure: Evidence From Animal and Human Studies. Alcohol Health and Research World, 18(1), 22–29.

de Mooij-van Malsen, J. G., Röhrdanz, N., Buschhoff, A.-S., Schiffelholz, T., Sigurdsson, T., & Wulff, P. (2023). Task-specific oscillatory synchronization of prefrontal cortex, nucleus reuniens, and hippocampus during working memory. iScience, 26(9), 107532. 10.1016/j.isci.2023.107532

Dobbing, J., & Sands, J. (1979). Comparative aspects of the brain growth spurt. Early Human Development, 3(1), 79–83. 10.1016/0378-3782(79)90022-7

Floresco, S. B., Seamans, J. K., & Phillips, A. G. (1997). Selective Roles for Hippocampal, Prefrontal Cortical, and Ventral Striatal Circuits in Radial-Arm Maze Tasks With or Without a Delay. The Journal of Neuroscience, 17(5), 1880. 10.1523/JNEUROSCI.17-05-01880.1997

George, A. E., Stout, J. J., & Griffin, A. L. (2023). Pausing and reorienting behaviors enhance the performance of a spatial working memory task. Behavioural Brain Research, 446, 114410. 10.1016/j.bbr.2023.114410

Grant, D. S. (1981). Intertrial interference in rat short-term memory. Journal of Experimental Psychology: Animal Behavior Processes, 7(3), 217–227. 10.1037/0097-7403.7.3.217

Griesbach, G. S., Hu, D., & Amsel, A. (1998). Effects of MK-801 on vicarious trial-and-error and reversal of olfactory discrimination learning in weanling rats. Behavioural Brain Research, 97(1–2), 29–38. 10.1016/s0166-4328(98)00015-1

Goodlett, C. R., Thomas, J. D., & West, J. R. (1991). Long-term deficits in cerebellar growth and rotarod performance of rats following “binge-like” alcohol exposure during the neonatal brain growth spurt. Neurotoxicology and Teratology, 13(1), 69–74. 10.1016/0892-0362(91)90029-V

Gursky, Z. H., Savage, L. M., & Klintsova, A. Y. (2019). Nucleus reuniens of the midline thalamus of a rat is specifically damaged after early postnatal alcohol exposure. NeuroReport, 30(10). https://journals.lww.com/neuroreport/Fulltext/2019/07010/Nucleus_reuniens_of_the_midline_thala mus_of_a_rat.10.aspx

Gursky, Z. H., Savage, L. M., & Klintsova, A. Y. (2021). Executive functioning-specific behavioral impairments in a rat model of human third trimester binge drinking implicate prefrontal-thalamo- hippocampal circuitry in Fetal Alcohol Spectrum Disorders. Behavioural Brain Research, 405, 113208. 10.1016/j.bbr.2021.113208

Gursky, Z. H., Spillman, E. C., & Klintsova, A. Y. (2020). Single-day Postnatal Alcohol Exposure Induces Apoptotic Cell Death and Causes long-term Neuron Loss in Rodent Thalamic Nucleus Reuniens. Neuroscience, 435, 124–134. 10.1016/j.neuroscience.2020.03.046

Hallock, H. L., Arreola, A. C., Shaw, C. L., & Griffin, A. L. (2013). Dissociable roles of the dorsal striatum and dorsal hippocampus in conditional discrimination and spatial alternation T-maze tasks. Neurobiology of Learning and Memory, 100, 108–116. 10.1016/j.nlm.2012.12.009

Hallock, H. L., Wang, A., & Griffin, A. L. (2016). Ventral Midline Thalamus Is Critical for Hippocampal–Prefrontal Synchrony and Spatial Working Memory. The Journal of Neuroscience, 36(32), 8372. 10.1523/JNEUROSCI.0991-16.2016

Hamilton, G. F., Hernandez, I. J., Krebs, C. P., Bucko, P. J., & Rhodes, J. S. (2017). Neonatal alcohol exposure reduces number of parvalbumin-positive interneurons in the medial prefrontal cortex and impairs passive avoidance acquisition in mice deficits not rescued from exercise. Neuroscience, 352, 52–63. 10.1016/j.neuroscience.2017.03.058

Hamilton, G. F., Whitcher, L. T., & Klintsova, A. Y. (2010). Postnatal binge-like alcohol exposure decreases dendritic complexity while increasing the density of mature spines in mPFC Layer II/III pyramidal neurons. Synapse, 64(2), 127–135. 10.1002/syn.20711

Hamre, K. M., & West, J. R. (1993). The Effects of the Timing of Ethanol Exposure during the Brain Growth Spurt on the Number of Cerebellar Purkinje and Granule Cell Nuclear Profiles. Alcoholism: Clinical and Experimental Research, 17(3), 610–622. 10.1111/j.1530-0277.1993.tb00808.x

Hoyme, H. E., Kalberg, W. O., Elliott, A. J., Blankenship, J., Buckley, D., Marais, A.-S., Manning, M. A., Robinson, L. K., Adam, M. P., Abdul-Rahman, O., Jewett, T., Coles, C. D., Chambers, C., Jones, K. L., Adnams, C. M., Shah, P. E., Riley, E. P., Charness, M. E., Warren, K. R., & May, P. A. (2016). Updated Clinical Guidelines for Diagnosing Fetal Alcohol Spectrum Disorders. Pediatrics, 138(2), e20154256. 10.1542/peds.2015-4256

Hu, D., & Amsel, A. (1995). A simple test of the vicarious trial-and-error hypothesis of hippocampal function. Proceedings of the National Academy of Sciences, 92(12), 5506–5509. 10.1073/pnas.92.12.5506

Ikonomidou, C., Bittigau, P., Ishimaru, M. J., Wozniak, D. F., Koch, C., Genz, K., Price, M. T., Stefovska, V., Hörster, F., Tenkova, T., Dikranian, K., & Olney, J. W. (2000). Ethanol-Induced Apoptotic Neurodegeneration and Fetal Alcohol Syndrome. Science, 287(5455), 1056–1060. 10.1126/science.287.5455.1056

Ito, H. T., Zhang, S.-J., Witter, M. P., Moser, E. I., & Moser, M.-B. (2015). A prefrontal–thalamo– hippocampal circuit for goal-directed spatial navigation. Nature, 522(7554), 50–55. 10.1038/nature14396

Janabi-Sharifi, F., Hayward, V., & Chen, C.-S. J. (2000). Discrete-time adaptive windowing for velocity estimation. IEEE Transactions on Control Systems Technology, 8(6), 1003–1009. 10.1109/87.880606

Jayachandran, M., Viena, T. D., Garcia, A., Veliz, A. V., Leyva, S., Roldan, V., Vertes, R. P., & Allen, T. A. (2023). Nucleus reuniens transiently synchronizes memory networks at beta frequencies. Nature Communications, 14(1), 4326. 10.1038/s41467-023-40044-z

Johnson, A., & Redish, A. D. (2007). Neural Ensembles in CA3 Transiently Encode Paths Forward of the Animal at a Decision Point. The Journal of Neuroscience, 27(45), 12176. 10.1523/JNEUROSCI.3761-07.2007

Jones, M. W., & Wilson, M. A. (2005). Theta Rhythms Coordinate Hippocampal–Prefrontal Interactions in a Spatial Memory Task. PLOS Biology, 3(12), e402. 10.1371/journal.pbio.0030402

Kay, K., Chung, J. E., Sosa, M., Schor, J. S., Karlsson, M. P., Larkin, M. C., Liu, D. F., & Frank, L. M. (2020). Constant Sub-second Cycling between Representations of Possible Futures in the Hippocampus. Cell, 180(3), 552–567.e25. 10.1016/j.cell.2020.01.014

Kidder, K. S., Miles, J. T., Baker, P. M., Hones, V. I., Gire, D. H., & Mizumori, S. J. Y. (2021). A selective role for the mPFC during choice and deliberation, but not spatial memory retention over short delays. Hippocampus, 31(7), 690–700. 10.1002/hipo.23306

Klintsova, A. Y., Cowell, R. M., Swain, R. A., Napper, R. M. A., Goodlett, C. R., & Greenough, W. T. (1998). Therapeutic effects of complex motor training on motor performance deficits induced by neonatal binge-like alcohol exposure in rats: I. Behavioral results. Brain Research, 800(1), 48–61. 10.1016/S0006-8993(98)00495-8

Lawrence, R. C., Otero, N. K. H., & Kelly, S. J. (2012). Selective effects of perinatal ethanol exposure in medial prefrontal cortex and nucleus accumbens. Neurotoxicology and Teratology, 34(1), 128–135. 10.1016/j.ntt.2011.08.002

Layfield, D. M., Patel, M., Hallock, H., & Griffin, A. L. (2015). Inactivation of the nucleus reuniens/rhomboid causes a delay-dependent impairment of spatial working memory. Neurobiology of Learning and Memory, 125, 163–167. 10.1016/j.nlm.2015.09.007

Livy, D. J., Miller, E. K., Maier, S. E., & West, J. R. (2003). Fetal alcohol exposure and temporal vulnerability: Effects of binge-like alcohol exposure on the developing rat hippocampus. Neurotoxicology and Teratology, 25(4), 447–458. 10.1016/s0892-0362(03)00030-8

Maharjan, D. M., Dai, Y. Y., Glantz, E. H., & Jadhav, S. P. (2018). Disruption of dorsal hippocampal – prefrontal interactions using chemogenetic inactivation impairs spatial learning. Neurobiology of Learning and Memory, 155, 351–360. 10.1016/j.nlm.2018.08.023

Mattson, S. N., Bernes, G. A., & Doyle, L. R. (2019). Fetal Alcohol Spectrum Disorders: A Review of the Neurobehavioral Deficits Associated With Prenatal Alcohol Exposure. Alcoholism: Clinical and Experimental Research, 43(6), 1046–1062. 10.1111/acer.14040

Miles, J. T., Mullins, G. L., & Mizumori, S. J. Y. (2024). Flexible decision-making is related to strategy learning, vicarious trial and error, and medial prefrontal rhythms during spatial set-shifting. *Learning & Memory (Cold Spring Harbor*, N.Y.), 31(7), a053911. 10.1101/lm.053911.123

Miller, E. K., & Cohen, J. D. (2001). An integrative theory of prefrontal cortex function. Annual Review of Neuroscience, 24, 167–202. 10.1146/annurev.neuro.24.1.167

Muenzinger, K. F. (1938). Vicarious Trial and Error at a Point of Choice: I. A General Survey of its Relation to Learning Efficiency. The Pedagogical Seminary and Journal of Genetic Psychology, 53(1), 75–86. 10.1080/08856559.1938.10533799

Murawski, N. J., Klintsova, A. Y., & Stanton, M. E. (2012). Neonatal alcohol exposure and the hippocampus in developing male rats: Effects on behaviorally induced CA1 c-Fos expression, CA1 pyramidal cell number, and contextual fear conditioning. Neuroscience, 206, 89–99. 10.1016/j.neuroscience.2012.01.006

O’Neill, P.-K., Gordon, J. A., & Sigurdsson, T. (2013). Theta oscillations in the medial prefrontal cortex are modulated by spatial working memory and synchronize with the hippocampus through its ventral subregion. The Journal of Neuroscience, 33(35), 14211–14224. 10.1523/JNEUROSCI.2378-13.2013

Otero, N. K. H., Thomas, J. D., Saski, C. A., Xia, X., & Kelly, S. J. (2012). Choline Supplementation and DNA Methylation in the Hippocampus and Prefrontal Cortex of Rats Exposed to Alcohol During Development. Alcohol: Clinical and Experimental Research, 36(10), 1701–1709. 10.1111/j.1530-0277.2012.01784.x

Papale, A. E., Stott, J. J., Powell, N. J., Regier, P. S., & Redish, A. D. (2012). Interactions between deliberation and delay-discounting in rats. *Cognitive, Affective*, & Behavioral Neuroscience, 12(3), 513–526. 10.3758/s13415-012-0097-7

Popova, S., Charness, M. E., Burd, L., Crawford, A., Hoyme, H. E., Mukherjee, R. A. S., Riley, E. P., & Elliott, E. J. (2023). Fetal alcohol spectrum disorders. Nature Reviews Disease Primers, 9(1), 11. 10.1038/s41572-023-00420-x

Rasmussen, C. (2005). Executive Functioning and Working Memory in Fetal Alcohol Spectrum Disorder. Alcoholism: Clinical and Experimental Research, 29(8), 1359–1367. 10.1097/01.alc.0000175040.91007.d0

Redish, A. D. (2016). Vicarious trial and error. Nature Reviews Neuroscience, 17(3), 147–159. 10.1038/nrn.2015.30

Sangiamo, D. T., Warren, M. R., & Neunuebel, J. P. (2020). Ultrasonic signals associated with different types of social behavior of mice. Nature Neuroscience, 23(3), 411–422. 10.1038/s41593-020-0584-z

Schmidt, B., Duin, A. A., & Redish, A. D. (2019). Disrupting the medial prefrontal cortex alters hippocampal sequences during deliberative decision making. Journal of Neurophysiology, 121(6), 1981–2000. 10.1152/jn.00793.2018

Schmidt, B., Papale, A., Redish, A. D., & Markus, E. J. (2013). Conflict between place and response navigation strategies: Effects on vicarious trial and error (VTE) behaviors. Learning & Memory, 20(3), 130–138. 10.1101/lm.028753.112

Steiner, A. P., & Redish, A. D. (2012). The Road Not Taken: Neural Correlates of Decision Making in Orbitofrontal Cortex. Frontiers in Neuroscience, 6. https://www.frontiersin.org/journals/neuroscience/articles/10.3389/fnins.2012.00131

Stout J. J., George A. E., Kim S., Hallock H. L., Griffin A. L. (2023). Using synchronized brain rhythms to bias memory-guided decisions. eLife 12:RP92033. 10.7554/eLife.92033.2

Stout, J. J., Hallock, H. L., George, A. E., Adiraju, S. S., & Griffin, A. L. (2022). The ventral midline thalamus coordinates prefrontal–hippocampal neural synchrony during vicarious trial and error. Scientific Reports, 12(1), 10940. 10.1038/s41598-022-14707-8

Tang, W., Shin, J. D., & Jadhav, S. P. (2021). Multiple time-scales of decision-making in the hippocampus and prefrontal cortex. eLife, 10, e66227. 10.7554/eLife.66227

Thomas, J. D., Wasserman, E. A., West, J. R., & Goodlett, C. R. (1996). Behavioral deficits induced by bingelike exposure to alcohol in neonatal rats: Importance of developmental timing and number of episodes. Developmental Psychobiology, 29(5), 433–452. 10.1002/(SICI)1098-2302(199607)29:5<433::AID-DEV3>3.0.CO;2-P

Thomas, J. D., Weinert, S. P., Sharif, S., & Riley, E. P. (1997). MK-801 Administration During Ethanol Withdrawal in Neonatal Rat Pups Attenuates Ethanol-Induced Behavioral Deficits. Alcohol: Clinical and Experimental Research, 21(7), 1218–1225. 10.1111/j.1530-0277.1997.tb04441.x

Tolman, E. C. (1939). Prediction of vicarious trial and error by means of the schematic sowbug. Psychological Review, 46(4), 318–336.

Tran, T. D., & Kelly, S. J. (2003). Critical periods for ethanol-induced cell loss in the hippocampal formation. Neurotoxicology and Teratology, 25(5), 519–528. 10.1016/S0892-0362(03)00074-6

Viena, T. D., Linley, S. B., & Vertes, R. P. (2018). Inactivation of nucleus reuniens impairs spatial working memory and behavioral flexibility in the rat. Hippocampus, 28(4), 297–311. 10.1002/hipo.22831

Wang, G.-W., & Cai, J.-X. (2006). Disconnection of the hippocampal–prefrontal cortical circuits impairs spatial working memory performance in rats. Behavioural Brain Research, 175(2), 329–336. 10.1016/j.bbr.2006.09.002

Wang, J. X., Cohen, N. J., & Voss, J. L. (2015). Covert rapid action-memory simulation (CRAMS): A hypothesis of hippocampal–prefrontal interactions for adaptive behavior. Memory and Decision Making, 117, 22–33. 10.1016/j.nlm.2014.04.003

Whitcher, L. T., & Klintsova, A. Y. (2008). Postnatal binge-like alcohol exposure reduces spine density without affecting dendritic morphology in rat mPFC. Synapse, 62(8), 566–573. 10.1002/syn.20532

Wozniak, D. F., Hartman, R. E., Boyle, M. P., Vogt, S. K., Brooks, A. R., Tenkova, T., Young, C., Olney, J. W., & Muglia, L. J. (2004). Apoptotic neurodegeneration induced by ethanol in neonatal mice is associated with profound learning/memory deficits in juveniles followed by progressive functional recovery in adults. Neurobiology of Disease, 17(3), 403–414. 10.1016/j.nbd.2004.08.006

